# Polyethylene glycol (PEG)-associated immune responses triggered by clinically relevant lipid nanoparticles in rats

**DOI:** 10.1101/2022.11.24.516986

**Authors:** Haiyang Wang, Yisha Wang, Changzheng Yuan, Xiao Xu, Wenbin Zhou, Yuhui Huang, Huan Lu, Yue Zheng, Gan Luo, Jia Shang, Meihua Sui

## Abstract

With the large-scale vaccination of lipid nanoparticles (LNP)-based COVID-19 mRNA vaccines, elucidating the potential polyethylene glycol (PEG)-associated immune responses triggered by clinically relevant LNP has become imminent. However, inconsistent findings were observed across very limited population-based studies. Herein we initiated a study using LNP carrier of Comirnaty^®^ as a representative, and simulated real-world clinical practice covering a series of time points and various doses correlated with approved LNP-delivered drugs in a rat model. We demonstrated the time- and dose-dependency of LNP-induced anti-PEG antibodies in rats. As a thymus-independent antigen, LNP unexpectedly induced isotype switch and immune memory, leading to rapid enhancement and longer lasting time of anti-PEG IgM and IgG upon re-injection in rats. Importantly, initial LNP injection accelerated the blood clearance of subsequent dosing in rats. These findings refine our understandings on LNP and possibly other PEG derivatives, and may promote optimization of related premarket guidelines and clinical protocols.

---

Lipid nanoparticles (LNP) composed of ionizable cationic lipid, cholesterol, 1,2-distearoyl-sn-glycero-3-phosphocholine (DSPC) and polyethylene glycol (PEG)-conjugated lipid, have attracted great attention due to unique advantages such as simple formulation, good biocompatibility and large payload^1^. Currently three LNP-delivered drugs have been marketed, including Patisiran (Onpattro^®^), mRNA vaccines BNT162b2 (Comirnaty^®^) and mRNA-1273 (Spikevax^®^)^2^. Modification of therapeutics with PEG has shown a number of advantages in several aspects^3^. Indeed, as the first vaccine using PEG as an excipient, its PEG-conjugated lipid composition (ALC-0159) plays critical roles in improving the stability and blood circulation of LNP, leading to the overwhelming success of Comirnaty^®4^. Although free PEG is poorly immunogenic, it has been recognized as a polyvalent hapten and may acquire immunogenic properties, *e.g.* inducing anti-PEG antibodies, upon conjugation with other materials such as proteins and nanocarriers^5–7^. Importantly, anti-PEG antibodies could form “antigen-antibody” complexes with newly administered PEGylated drugs, while subsequent clearance of these complexes by macrophage may lead to biodistribution/pharmacokinetic changes and reduced efficacy of PEGylated drugs^5,6^. Moreover, “antigen-antibody” complexes may induce severe side effects including hypersensitivity reactions, although the underlying mechanisms have not been fully clarified^6,8^. Interestingly, a proportion of individuals who never received PEGylated drugs have anti-PEG antibodies possibly due to environmental exposure^9,10^.

With the large-scale vaccination of mRNA vaccines, elucidating the potential PEG-associated immune responses triggered by clinically relevant LNP has become urgent^11^. However, there are only five related literatures, all of which are recent clinical observations^12–16^. It is noteworthy that several limitations are existed in these studies, *e.g.* small population size^13, 15^, large person-to-person variability of pre-existing anti-PEG antibodies^13, 14^, age- and gender-related influences^12–14, 16^, unavoidable exposure to PEG-containing substances other than LNP^12–16^, deviation of sampling time points^12–16^, and mixed use of different LNP-delivered drugs^14^. Currently, there is no consistent conclusion regarding any characteristic of initial and/or repeated injection of mRNA vaccines in inducing PEG-specific antibodies. Moreover, the amount of mPEG_2000_ contained in each injection varies significantly among three approved LNP-delivered drugs^17–19^. For example, mPEG_2000_ contained in each Onpattro^®^ injection is as high as 262 times of Comirnaty^®17^, raising our concern on the potential impact of mPEG_2000_ exposure amount on induction of PEG-associated immune responses. Furthermore, as the first two vaccines using LNP as carriers, the pharmacokinetics of Comirnaty^®^ and Spikevax^®^ might differ from previously approved intramuscular vaccines, considering that the *in vivo* procedures of mRNA vaccines are mainly determined by their LNP carriers^18^. However, pharmacokinetic data is not available for either Comirnaty^®^ or Spikevax^®^, as these data are not regularly required by WHO for market approval of intramuscular vaccines^20^.

To clarify the cause-and-effect relationship of clinically relevant LNP in inducing PEG-associated immune responses, we synthesized the LNP of Comirnaty^®^ (most widely used clinically relevant LNP) as a representative. A Wistar rat model with excellent quality control was established to eliminate undesired interferences existed in previous studies. Through simulating the clinical practice of Comirnaty^®^, including two intramuscular injections with a 21-day interval, and delicately designing clinically relevant doses covering the whole range of mPEG_2000_ amount contained in a single injection of approved LNP-delivered drugs, the PEG-associated immune responses triggered by LNP including potential impact on pharmacokinetic changes were carefully investigated.

## Results

### Synthesis and physiochemical characterization of PEGylated LNP, DiR-LNP and DiR-LU@LNP

LNP, DiR-LNP (LNP labeled with DiR) and DiR-LU@LNP (LNP loaded with firefly luciferase mRNA and labeled with DiR) were prepared by mixing the ethanol phase containing ALC-0315, DSPC, cholesterol and ALC-0159 (with or without DiR) and the aqueous phase containing citrate buffer (with or without firefly luciferase mRNA) through a microfluidic mixing device (**Fig. 1a-1c**). When examined with Cryo-TEM, LNP and DiR-LNP were characterized as hollow spheres, while DiR-LU@LNP exhibited a typical electron-dense core structure containing mRNA (**Fig. 1c**). Further characterization with DLS showed that the Z-average/PDI/Zeta potential of LNP, DiR-LNP and DiR-LU@LNP were 110.400 ± 3.466 nm/0.203 ± 0.012/16.733 ± 0.451 mV, 113.067 ± 2.139 nm/0.183 ± 0.013/7.257 ± 0.168 mV and 101.367 ± 2.593 nm/0.197 ± 0.015/-5.943 ± 0.129 mV, respectively (**Fig. 1d and 1e**; **Supplementary Table 2**). These data demonstrate that three LNP formulations have optimal particle diameter, highly monodisperse particle-size distribution and weak surface charge. Additional evaluation in the presence of 10% rat serum mimicking blood circulation showed that LNP, DiR-LNP and DiR-LU@LNP have relatively stable particle sizes and stay monodisperse (**Supplementary** Fig. 1a-1c**; Supplementary** Figs. 2-4). Moreover, standard curves of phospholipid (DSPC) contained in three LNP formulations were respectively drawn, as the phospholipid component is commonly used for quantifying the whole LNP^21,22^. Subsequently, clinically relevant doses of LNP including low dose (L-LNP, 0.009 mg phospholipids/kg), middle dose (M-LNP, 0.342 mg phospholipids/kg) and high dose (H-LNP, 2.358 mg phospholipids/kg) were calculated based on the corresponding equations (**Supplementary** Fig. 1d-f; see **Methods**).

**Fig. 1.**
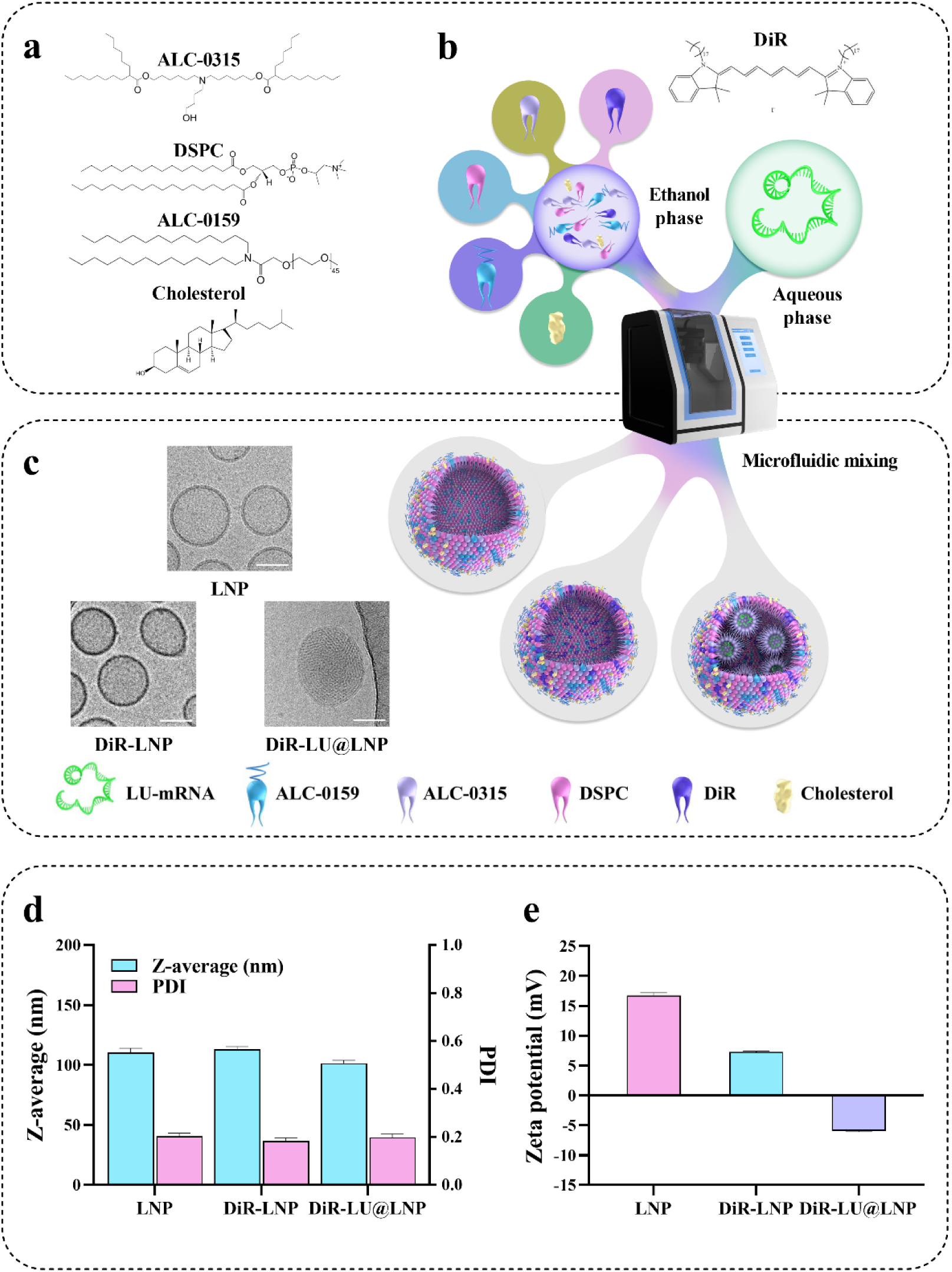
Preparation and characterization of LNP, DiR-LNP and DiR-LU@LNP. (**a**) Chemical structures of lipid compositions in LNP carrier of COVID-19 vaccine Comirnaty^®^. (**b**) Schematic illustration of the synthesis of LNP, DiR-LNP and DiR-LU@LNP. Briefly, the ethanol phase was combined with the aqueous phase at a flow rate ratio of 1: 3 (ethanol: aqueous) through a microfluidic mixing device. (**c**) Representative cryogenic transmission electron microscopy (Cryo-TEM) images of LNP, DiR-LNP and DiR-LU@LNP. Scale bar: 50 nm. (**d**) Hydrodynamic size (Z-average) and polydispersity index (PDI) of LNP, DiR-LNP and DiR-LU@LNP measured by DLS. (**e**) Zeta potential of LNP, DiR-LNP and DiR-LU@LNP measured by DLS. Data in **d** and **e** were presented as “mean ± standard deviation” (n=3). LNP, lipid nanoparticles; DLS, dynamic light scattering; LU-mRNA: luciferase mRNA.

### Time- and dose-dependent induction of anti-PEG IgM by PEGylated LNP

After intramuscular injection of LNP at above-mentioned three doses on Day 0 and Day 21, rat serum samples were respectively collected at 12 designated time points between Day 0 and Day 49, and examined for the presence and level of anti-PEG IgM (**Fig. 2a**). Our data showed that serum anti-PEG IgM was initially detected in L-LNP group on Day 3. Although anti-PEG IgM was undetectable until Day 5 in M-LNP and H-LNP groups, both doses induced significantly higher levels of anti-PEG IgM than that induced by L-LNP. Moreover, L-LNP induced anti-PEG IgM only detectable on Day 3 and Day 5 during the first injection cycle (Day 0∼21), while M-LNP and H-LNP induced more persistent and higher levels of anti-PEG IgM detectable throughout Day 5∼21, suggesting a time- and dose-dependent induction of anti-PEG IgM after an initial injection of LNP. Impressively, anti-PEG IgM was detected at more time points for all three doses after repeated injection compared with that during the first injection cycle. Particularly, M-LNP and H-LNP constantly induced anti-PEG IgM throughout the whole second injection cycle and extension period (Day 21∼49). Meanwhile, anti-PEG IgM levels induced by three doses exhibited a consistent ranking (H-LNP > M-LNP > L-LNP) at nearly all detectable time points (**Fig. 2b**). In addition, the intra-assay precision/Coefficient of Variation (CV%) of anti-PEG IgM standards and serum samples were 3.884 ± 3.046% and 5.237 ± 6.192%, respectively (**Fig. 2c**), and the inter-assay precision/Coefficient of Variation (CV%) of anti-PEG IgM standards and goodness of fit/coefficient of determination (*R*^2^) of the standard curve were 20.983 ± 15.511% and 0.992 ± 0.004, respectively (**Fig. 2c**; **Supplementary** Fig. 5). These data demonstrate the good quality control of ELISA (see **Methods** for acceptance criteria^23,24^). Overall, these data provided strong evidence for the dose- and time-dependent induction of anti-PEG IgM.

**Fig. 2.**
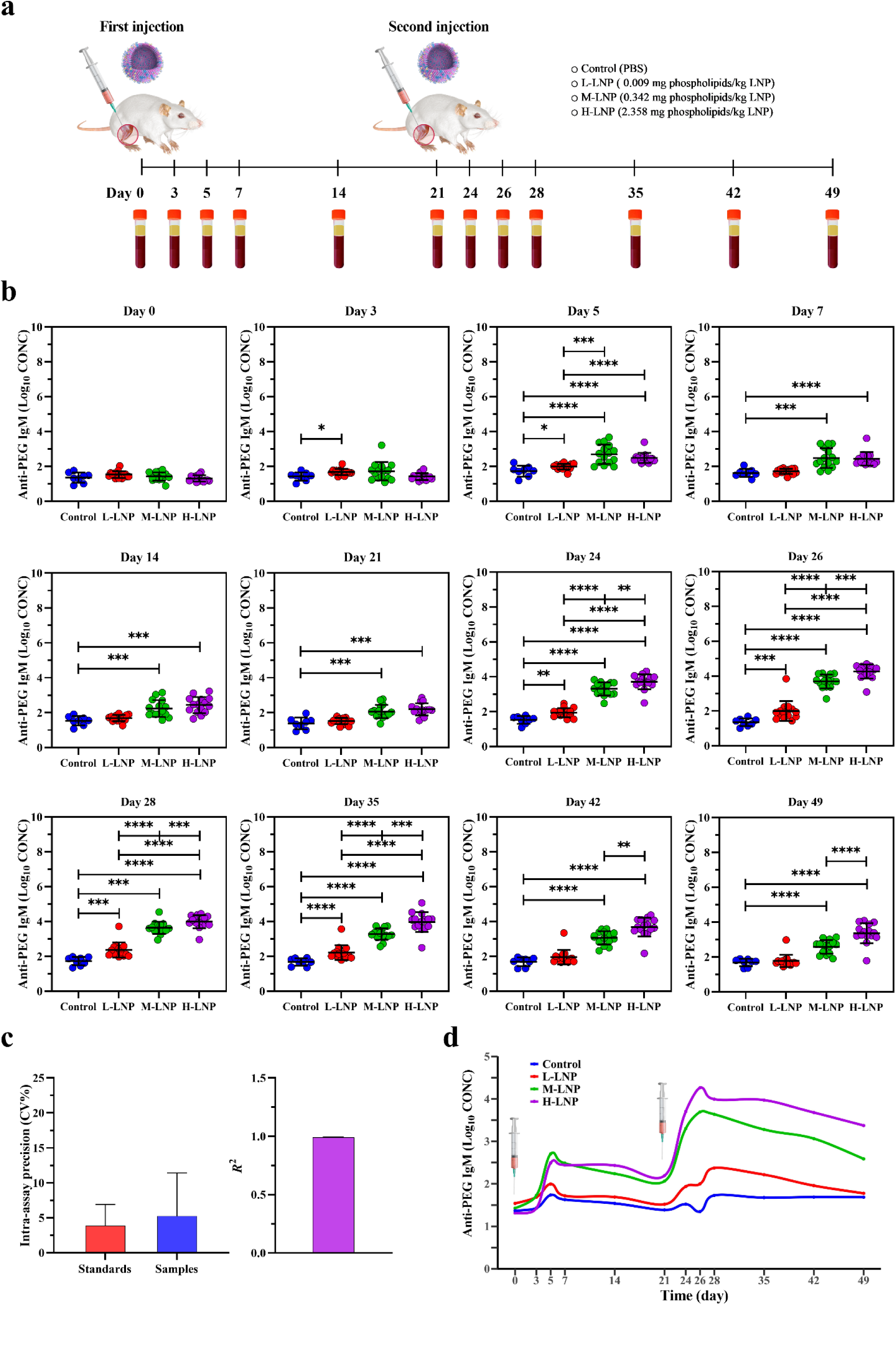
Experimental design and evaluation of anti-PEG IgM production in rat. (**a**) Schematic illustration of the experimental protocols. Wistar rats were injected intramuscularly with 0.009 (L-LNP group), 0.342 (M-LNP group) or 2.358 (H-LNP group) mg phospholipids/kg LNP on Day 0 and Day 21, respectively. Rats in the Control group were injected with PBS. Serum samples were collected at the indicated time points (Day 0, 3, 5, 7, 14, 21, 24, 26, 28, 35, 42 and 49) for further evaluation of the presence and level of anti-PEG antibodies with ELISA. (**b**) Quantitative analysis of anti-PEG IgM (Log_10_ CONC) (log_10_-transformed concentration of anti-PEG IgM) induced by LNP in rat serum. Data were presented as “mean ± standard deviation”, with n = 8 for Control group and n = 15 for all LNP-treated groups. Differences in anti-PEG IgM (Log_10_ CONC) among various groups were analyzed using Mann-Whitney U test, with *P* values adjusted for FDR (false discovery rate). Peak levels of anti-PEG IgM (Log_10_ CONC) induced during the initial and second injection cycle were as follows: L-LNP, 1.996 on Day 5 and 2.374 on Day 28; M-LNP, 2.704 on Day 5 and 3.692 on Day 26; H-LNP, 2.492 on Day 5 and 4.262 on Day 26. *, *P* < 0.05; **, *P* < 0.01; ***, *P* < 0.001; ****, *P* < 0.0001. (**c**) Representative quality control data of ELISA for detecting anti-PEG IgM. The left image shows the intra-assay precision/Coefficient of Variation (CV%) of anti-PEG IgM standards (CV% = 3.884 ± 3.046%) and serum samples (CV% = 5.237 ± 6.192%), and the right image shows the goodness of fit/coefficient of determination (*R*^2^ *=* 0.992 ± 0.004) of the standard curve (see **Methods** for acceptance criteria). Data were presented as “mean ± standard deviation”, with n = 56 for intra-assay precision (CV%) of anti-PEG IgM standards, n = 636 for intra-assay precision (CV%) of serum samples and n = 8 for *R*^2^. (**d**) Time-course of anti-PEG IgM induced by PEGylated LNP. The changing curves of mean anti-PEG IgM (Log_10_ CONC) levels over time were fitted by the R package called “ggalt”.

Additionally, anti-PEG IgM production exhibited different time-course profiles across different doses (**Fig. 2d**; all *P* < 0.05 in profile analysis). Further linear mixed model (LMM) analysis was conducted to evaluate changes over time and differences across groups regarding anti-PEG IgM production (**Supplementary Table 3**). Our data showed that β for “Group”, which represents mean differences on antibody level among various groups at all time points, exhibited statistical significance between Control *vs* M-LNP, Control *vs* H-LNP, and L-LNP *vs* M-LNP, respectively. Significant differences were also detected with β for “Time” and “Time^2^”, both of which representing the change rate in antibody level over time. Regarding β for “Group*Time” representing mean differences in the change rate in antibody level over time among various groups, we found that compared with the Control group, M-LNP and H-LNP groups showed faster rate of anti-PEG IgM production, with H-LNP group exhibited the fastest rate among all the groups, which further demonstrated the above-mentioned dose- and time-dependency. Moreover, consistent with the longer lasting period and higher level of anti-PEG IgM induced by repeated LNP injection, LMM analysis revealed the significant difference on antibody level between two separate injections (β for “Second Injection”: *P* < 0.0001 *vs* First Injection).

### Time- and dose-dependent induction of anti-PEG IgG by PEGylated LNP

Serum samples collected at above-mentioned time points were further examined for anti-PEG IgG (**Fig. 3a**). Interestingly, different from anti-PEG IgM production, no anti-PEG IgG was detected throughout the first injection cycle in all experimental groups. These data suggest that an initial single injection of LNP, at a broad range of doses, could not induce anti-PEG IgG in rats. However, although anti-PEG IgG was still undetectable after the second L-LNP injection, it was clearly induced by the repeated injection of M-LNP and H-LNP, and persisted at all later time points (Day 24∼49). Similar to the dose-dependency of anti-PEG IgM, anti-PEG IgG levels induced by H-LNP were significantly higher than corresponding levels induced by M-LNP at all detectable time points, with peak levels achieved on Day 26 for both M-LNP and H-LNP groups. Moreover, the intra-assay precision/Coefficient of Variation (CV%) of anti-PEG IgG standards and serum samples were 4.897 ± 5.549% and 8.546 ± 12.211%, respectively, and the inter-assay precision/Coefficient of Variation (CV%) of anti-PEG IgG standards and goodness of fit/coefficient of determination (*R*^2^) were 24.896 ± 10.071% and 0.999 ± 0.001, respectively (**Fig. 3b**; **Supplementary** Fig. 6), all of which demonstrating the good quality control of ELISA (see **Methods** for acceptance criteria^23,24^).

**Fig. 3.**
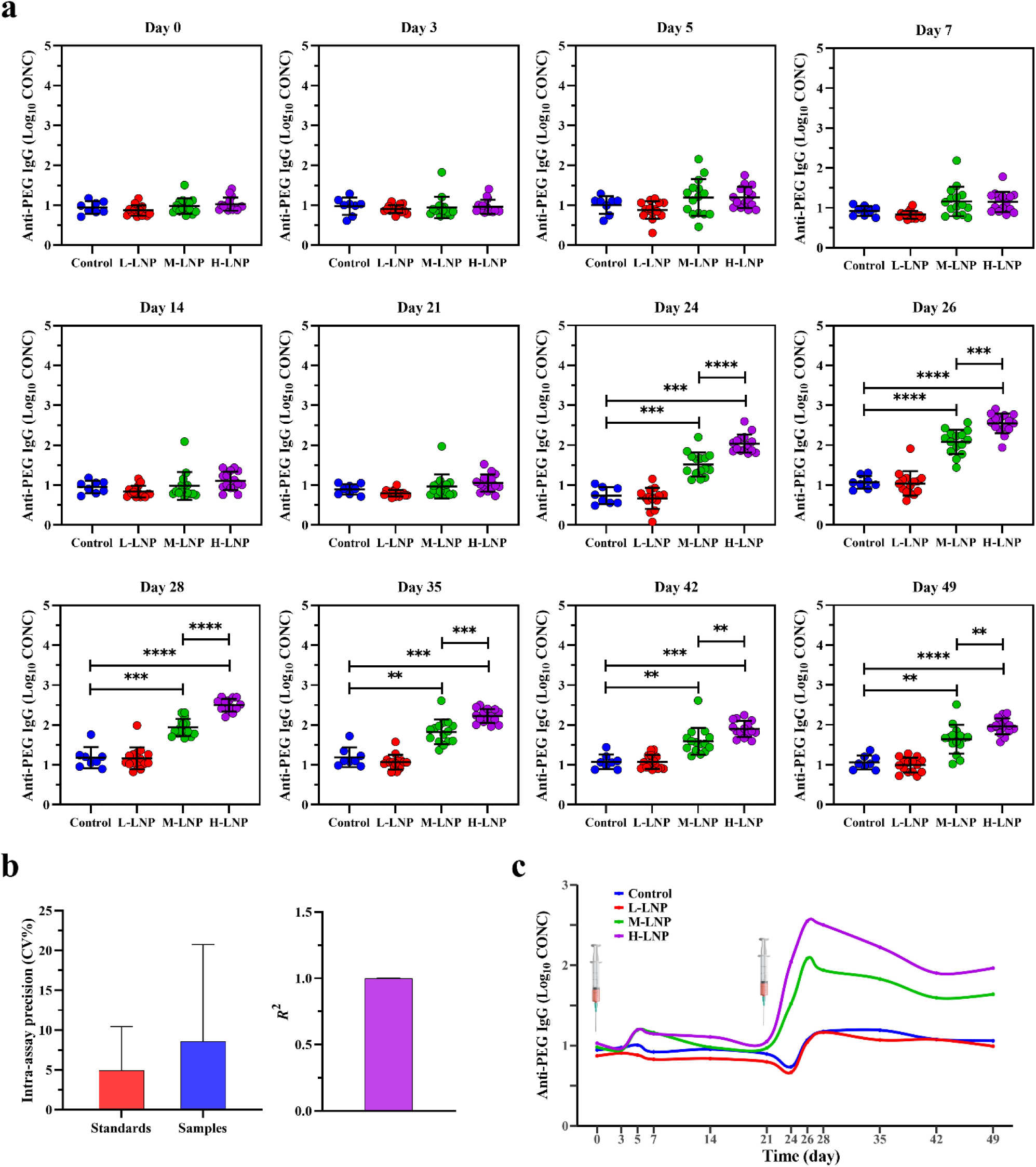
Evaluation of anti-PEG IgG production in rat. (**a**) Quantitative analysis of anti-PEG IgG (Log_10_ CONC) (log_10_-transformed concentration of anti-PEG IgG) induced by LNP in rat serum. Data were presented as “mean ± standard deviation”, with n = 8 for Control group and n = 15 for all LNP-treated groups. Differences in anti-PEG IgG (Log_10_ CONC) among various groups were analyzed using Mann-Whitney U test, with *P* values adjusted for FDR (false discovery rate). Peak levels of anti-PEG IgG (Log_10_ CONC) induced during the second injection cycle were as follows: M-LNP, 2.083 on Day 26; H-LNP, 2.547 on Day 26. *, *P* < 0.05; **, *P* < 0.01; ***, *P* < 0.001; ****, *P* < 0.0001. (**b**) Representative quality control data of ELISA for detecting anti-PEG IgG. The left image shows the intra-assay precision/Coefficient of Variation (CV%) of anti-PEG IgG standards (CV% = 4.897 ± 5.549%) and serum samples (CV% = 8.546 ± 12.211%), and the right image shows the goodness of fit/coefficient of determination (*R*^2^ *=* 0.999 ± 0.001) of the standard curve (see **Methods** for acceptance criteria). Data were presented as “mean ± standard deviation”, with n = 56 for intra-assay precision (CV%) of anti-PEG IgM standards, n = 636 for intra-assay precision (CV%) of serum samples and n = 8 for *R*^2^. (**c**) Time-course of anti-PEG IgG induced by PEGylated LNP. The changing curves of mean anti-PEG IgG (Log_10_ CONC) levels over time were fitted by the R package called “ggalt”.

As depicted in **Fig. 3c**, Control and L-LNP groups had similar time-course profile in anti-PEG IgG production (*P* > 0.05 in profile analysis), whereas every other two groups exhibited different profiles (all *P* < 0.05 in profile analysis). Further LMM analysis was conducted to evaluate the changes of anti-PEG IgG over time and differences across groups (**Supplementary Table 4**). β for “Group” exhibited statistical significance between M-LNP *vs* L-LNP and H-LNP *vs* L-LNP. Similar to anti-PEG IgM induction, significant differences were detected with β for “Time” and “Time^2^”. Meanwhile, both M-LNP and H-LNP groups had faster rate of anti-PEG IgG production compared with the Control group regarding β for “Group*Time”, with H-LNP group exhibited the fastest rate among all experimental groups. These data further demonstrate dose- and time-dependent induction of anti-PEG IgG by LNP. Correspondingly, LMM analysis confirmed the significant difference on anti-PEG IgG level between the two separate injections (β for “Second Injection”: *P* < 0.0001 *vs* First Injection).

### Enhanced production of anti-PEG antibodies by previous exposure to PEGylated LNP

To quantify the potential influence of initial/previous exposure to LNP on the production of anti-PEG antibodies after repeated LNP injection, increased anti-PEG IgM (▴Anti-PEG IgM (Log_10_ CONC)) and increased anti-PEG IgG (▴Anti-PEG IgG (Log_10_ CONC)) were respectively calculated by subtracting log10-transformed anti-PEG antibody concentration after first injection from that after the second injection for all tested doses and time points (▴Day 0, 3, 5, 7, 14 and 21). Our data showed that although ▴Anti-PEG IgM (Log_10_ CONC) was undetectable until ▴Day 5 in L-LNP group, significantly increased anti-PEG IgM production was observed at all time points in M-LNP and H-LNP groups. Moreover, two sequential injections of L-LNP induced transient ▴Anti-PEG IgM (Log_10_ CONC) detectable on ▴Day 5∼14, while those of M-LNP and L-LNP induced more persistent and higher level of ▴Anti-PEG IgM (Log_10_ CONC) detectable throughout ▴Day 3∼21 (**Fig. 4a**). Further profile analysis revealed that time-course of▴Anti-PEG IgM (Log_10_ CONC) between every two groups exhibited different profiles (**Fig. 4b**; all *P* < 0.05 in profile analysis). Coincidentally, LMM analysis demonstrated statistical significance on ▴Anti-PEG IgM (Log_10_ CONC) changes over 6 time points (except for H-LNP *vs* M-LNP) and differences across 3 doses (**Supplementary Table 5**). For instance, ▴Anti-PEG IgM (Log_10_ CONC) ranking from low to high was that respectively induced by L-LNP, M-LNP and H-LNP at all detectable time points regarding β for “Group”, demonstrating the dose dependency of ▴Anti-PEG IgM (Log_10_ CONC). Additionally, change rate in ▴Anti-PEG IgM (Log_10_ CONC) exhibited significant differences over 6 time points, with *P* < 0.0001 for both “Time” and “Time^2^”. Regarding β for “Group*Time”, M-LNP and H-LNP groups had faster change rate in ▴Anti-PEG IgM (Log_10_ CONC) than L-LNP group. These data have provided additional evidence for dose- and time-dependency of ▴Anti-PEG IgM (Log_10_ CONC) induced by two sequential injections of LNP.

**Fig. 4.**
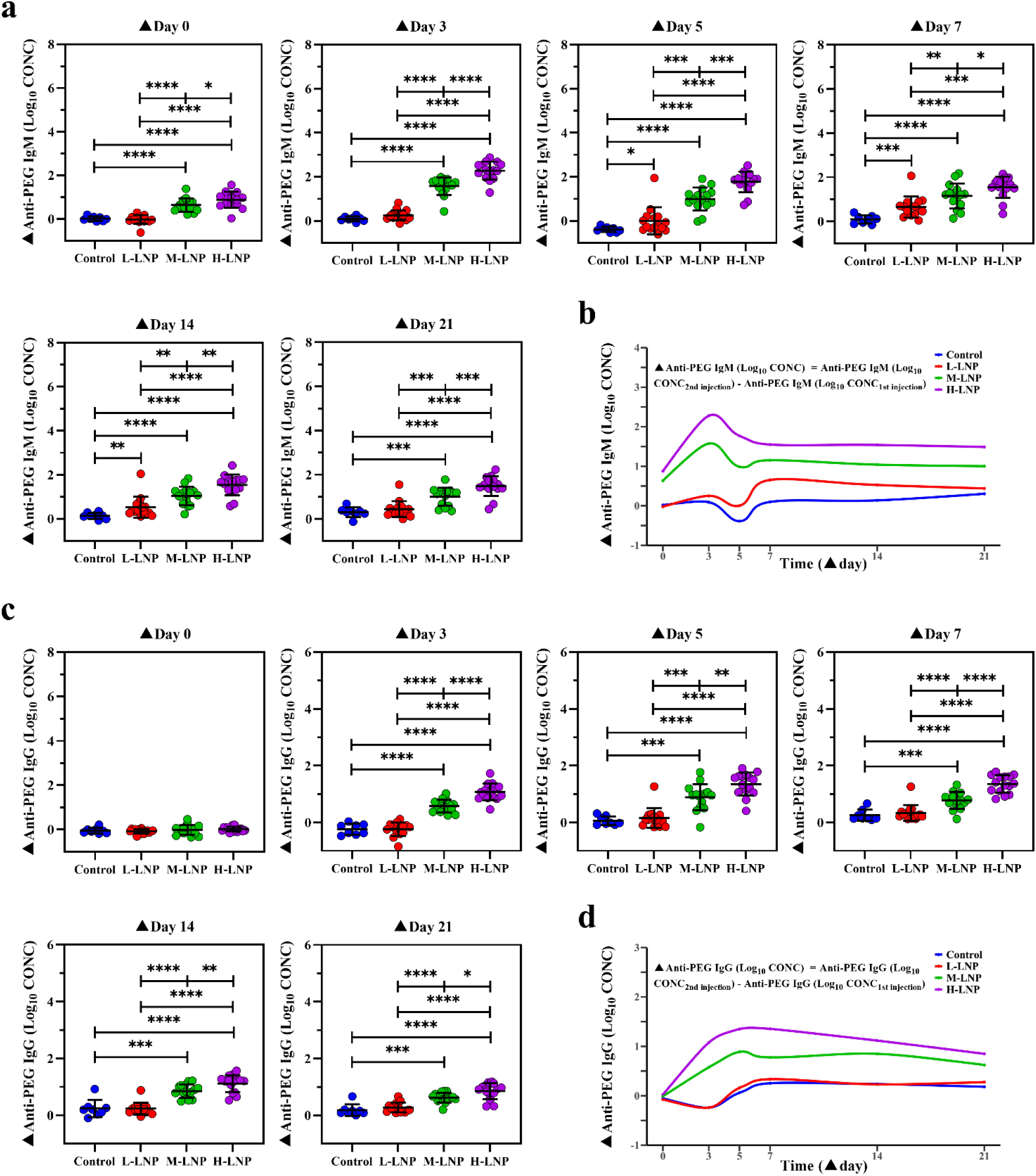
Enhanced production of anti-PEG antibodies in rat by repeated administration with PEGylated LNP. (**a**) Enhanced anti-PEG IgM production induced by repeated LNP injection. ▴Anti-PEG IgM (Log_10_ CONC) means Anti-PEG IgM (Log_10_ CONC_2nd injection_) (log_10_-transformed concentration of anti-PEG IgM induced during the second injection cycle) subtracted corresponding Anti-PEG IgM (Log_10_ CONC_1st injection_) (log_10_-transformed concentrations of anti-PEG IgM induced during the first injection cycle). Peak levels of ▴Anti-PEG IgM (Log_10_ CONC) induced by different LNP doses were 0.654 ± 0.471 for L-LNP (▴Day 7), 1.574 ± 0.399 for M-LNP (▴Day 3) and 2.277 ± 0.410 for H-LNP (Day 3), respectively, with significant difference among three groups (*P* < 0.001 for M-LNP *vs* L-LNP; *P* < 0.0001 for H-LNP vs L-LNP; *P*<0.0001 for H-LNP *vs* M-LNP). (**b**) Time-course of enhanced anti-PEG IgM induced by repeated injection of LNP. (c) Enhanced anti-PEG IgG production induced by repeated injection of LNP. ▴Anti-PEG IgG (Log_10_ CONC) means Anti-PEG IgG (Log_10_ CONC_2nd injection_) (log10-transformed concentration of anti-PEG IgG induced during the second injection cycle) subtracted corresponding Anti-PEG IgG (Log_10_ CONC_1st injection_) (log_10_-transformed concentrations of anti-PEG IgG induced during the first injection cycle). Peak levels of ▴Anti-PEG IgG (Log_10_ CONC) induced by different LNP doses were 0.888 ± 0.459 for M-LNP (▴Day 5) and 1.354 ± 0.308 for H-LNP (▴Day 7), respectively, with significant difference between these two groups (*P* < 0.01). (**d**) Time-course of enhanced anti-PEG IgG induced by repeated injection of LNP. In figure **a** and **c**, data were presented as “mean ± standard deviation”, with n = 8 for Control group and n = 15 for all LNP-treated groups. Differences in ▴Anti-PEG IgM (Log_10_ CONC) or ▴Anti-PEG IgG (Log_10_ CONC) among various groups were analyzed using Mann-Whitney U test, with *P* values adjusted for FDR (false discovery rate). *, *P* < 0.05; **, *P* < 0.01; ***, *P* < 0.001; ****, *P* < 0.0001. In Figure **b** and **d**, changing curves of average level of ▴Anti-PEG IgM (Log_10_ CONC) or ▴Anti-PEG IgG (Log_10_ CONC) over time for various doses were fitted by the R package called “ggalt”.

As anti-PEG IgG was absent throughout the first injection cycle and in all L-LNP-treated groups, ▴Anti-PEG IgG (Log_10_ CONC) was detected on ▴Day 3∼21 in M-LNP and H-LNP groups (**Fig. 4c**). Our data showed that the time-course profiles of ▴Anti-PEG IgG (Log_10_ CONC) exhibited significant difference among all other experimental groups (all *P* < 0.05 in profile analysis), except that no difference between Control and L-LNP groups was found (*P* > 0.05 in profile analysis). Corresponding LMM analysis further confirmed these findings (**Supplementary Table 6**), as statistical significances were obtained for ▴Anti-PEG IgG (Log_10_ CONC) among Control/L-LNP, M-LNP and H-LNP groups regarding β for “Group”. Together with the significant differences for “Time” and “Time^2^” on change rate over 6 time points, our data clearly demonstrated that similar to anti-PEG IgM, an initial/previous injection of LNP dose- and time-dependently boosted the generation of anti-PEG IgG after the repeated injection.

### Dose-dependent biodistribution of PEGylated LNP administered at clinically relevant doses

By using a fluorescence and bioluminescence double-labeling strategy, the biodistribution of LNP was determined in rats treated with DiR-LU@LNP simulating clinical practice (**Fig. 5a**). Consistent with the preclinical biodistribution data published in Assessment Report of Comirnaty^®^ issued by the European Medicines Agency, weak bioluminescence signal of luciferase was detected in muscle at injection site and liver (**Supplementary** Fig. 7), demonstrating that DiR-LU@LNP drained into the liver and delivered active luciferase mRNA. As DiR fluorescence exhibited significantly higher sensitivity than luciferase bioluminescence (**Fig. 5b**; **Supplementary** Fig. 7), LNP biodistribution was further analyzed based on DiR fluorescence. Our data showed that 6 hours after both the first and second injections, DiR fluorescence was only detectable in muscle at the injection site in L-LNP group. Upon increase of LNP dose, the fluorescent signal was significantly enhanced and detected in more organs/tissues (muscle at the injection site, liver and lung in M-LNP group; muscle at the injection site, liver, lung, spleen and draining lymph node in H-LNP group). Further analysis indicated that the total radiant efficiency from liver, lung, spleen and heart exhibited statistical significance between Control *vs* M-LNP, Control *vs* H-LNP, L-LNP *vs* M-LNP, L-LNP *vs* H-LNP, and M-LNP *vs* H-LNP after both the first and second injections. These findings demonstrate a dose-dependent biodistribution of LNP, with preferential accumulation in reticuloendothelial system after entering the blood circulation *via* intramuscular injection (**Fig. 5b and 5c**).

**Fig. 5.**
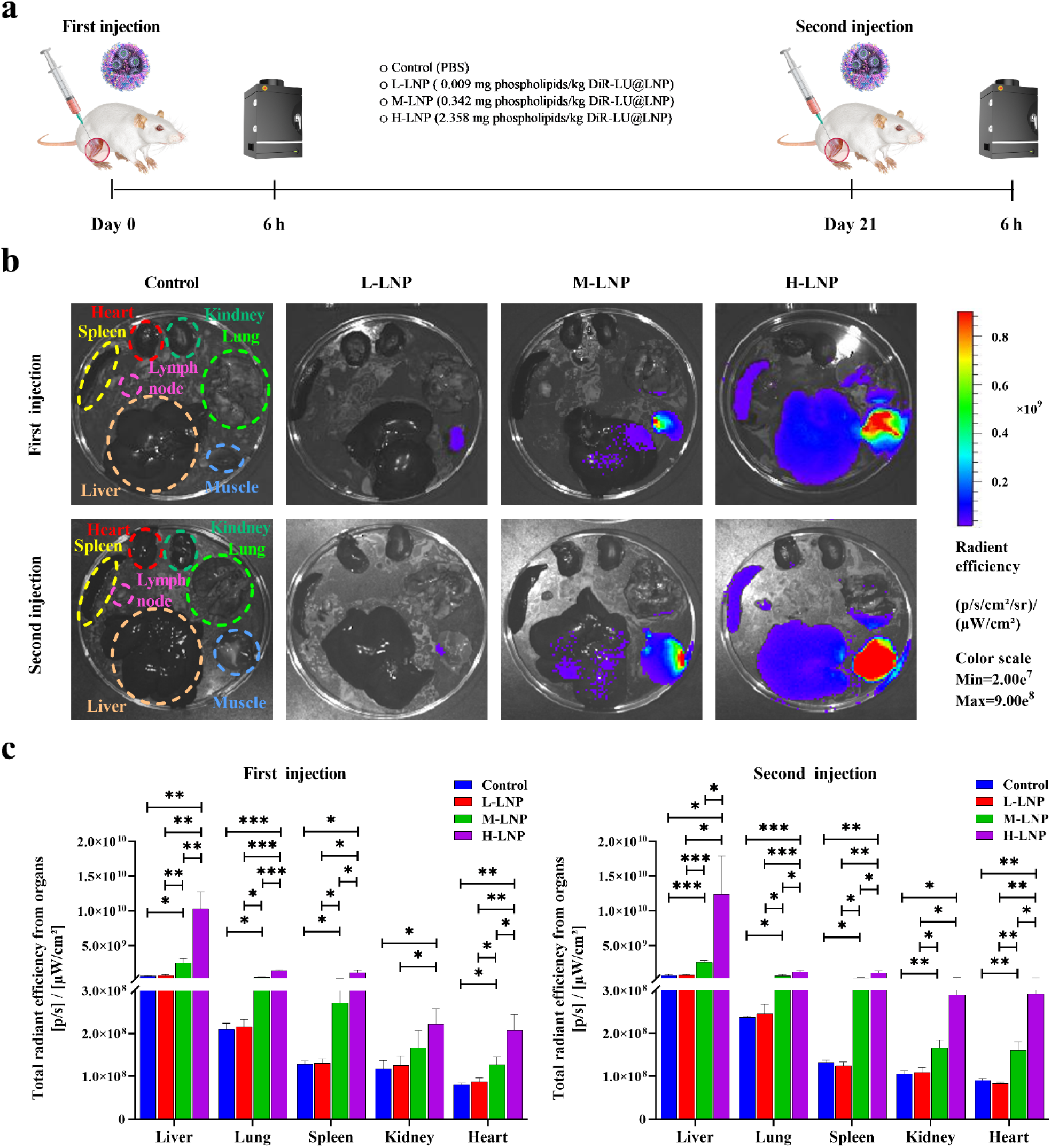
Experimental design and biodistribution of PEGylated LNP in representative organs of rat. (**a**) Schematic illustration of the experimental protocols. Wistar rats were injected intramuscularly with 0.009 (L-LNP group), 0.342 (M-LNP group) or 2.358 (H-LNP group) mg phospholipids/kg DiR-LU@LNP on Day 0 and Day 21, respectively. Rats in the Control group were injected with PBS. Six hours after each injection, three rats from each experimental group were sacrificed and immediately dissected. Major organs including heart, liver, spleen, lung, kidneys and draining lymph node, and muscle at the injection site were collected for fluorescence imaging with IVIS Spectrum imaging system. (**b**) Representative fluorescence images of major organs and muscle tissues isolated from rats 6 hours after the first and second injection of DiR-LU@LNP. (**c**) Total radiant efficiency of major organs determined 6 hours after the first and second injection of DiR-LU@LNP. Data were presented as “mean ± standard deviation” (n = 3). Differences in total radiation efficiency induced by three doses were analyzed using multiple unpaired *t* tests with correction for multiple testing. *, *P* < 0.05; **, *P* < 0.01; ***, *P* < 0.001; ****, *P* < 0.0001.

### Accelerated blood clearance induced by repeated injection of PEGylated LNP administered at clinically relevant dose

To explore whether previous exposure would alter the pharmacokinetic of repeatedly or newly injected LNP, rats were injected with DiR-LNP at above-mentioned doses and schedule, followed by collection of serum samples at 8 time points after each injection and measurement of DiR fluorescence (**Fig. 6a**). Our data indicate that LNP-associated DiR fluorescence was undetectable in L-LNP group at all time points after both injections, suggesting a low level of LNP in blood circulation after administration of low dose DiR-LNP. As expected, DiR fluorescence was significantly increased in serums isolated from M-LNP (at 6 and 10 hours) and H-LNP (at 7 sequential time points between 30 minutes and 48 hours) groups after the initial injection of DiR-LNP. However, compared with the first injection, DiR fluorescence was detected at less time points in M-LNP (only 6 hours) and H-LNP (4 sequential time points between 6 and 48 hours) groups after the second injection of DiR-LNP, indicating faster blood clearance and reduced serum level of LNP upon repeated administrations (**Fig. 6b**). Indeed, as depicted in **Fig. 6c and 6d**, DiR fluorescence was significantly decreased at 30 minutes, 1 hour and 48 hours after repeated injection of high dose DiR-LNP compared with that after the initial injection. These data demonstrate for the first time an accelerated blood clearance (ABC) phenomenon of clinically relevant LNP.

**Fig. 6.**
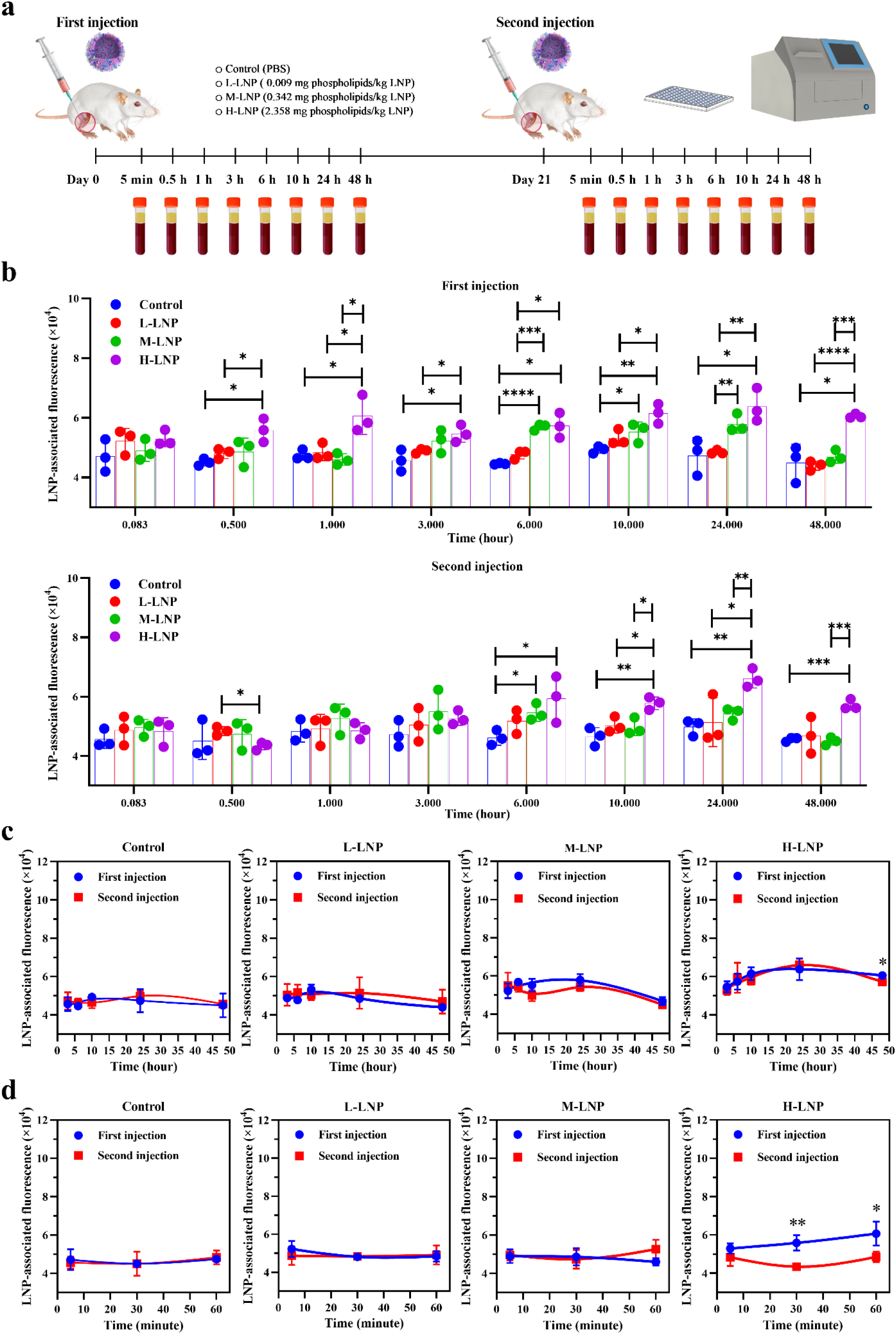
Experimental design and blood clearance of PEGylated LNP in rats. (**a**) Schematic illustration of the experimental protocols. Wistar rats were injected intramuscularly with 0.009 (L-LNP group), 0.342 (M-LNP group) or 2.358 (H-LNP group) mg phospholipids/kg DiR-LNP on Day 0 and Day 21, respectively. Rats in the Control group were injected with PBS. Serum samples were collected at the indicated 8 time points (5 minutes, 30 minutes, 1 hour, 3 hours, 6 hours, 10 hours, 24 and 48 hours) after each injection of DiR-LNP, followed by determination of LNP-associated fluorescence with Spectramax ID5 fluorescent spectrometry. (**b**) LNP-associated fluorescence was presented as “mean ± standard deviation” (n = 3) for each group, with differences among various groups after each injection analyzed using the multiple unpaired *t* test, with *P* values adjusted for FDR (false discovery rate). *, *P* < 0.05; **, *P* < 0.01; ***, *P* < 0.001; ****, *P* < 0.0001. (**c**) Blood clearance profile of DiR-LNP in rats based on LNP-associated fluorescence obtained at 3 hours, 6 hours, 10 hours, 24 and 48 hours, with fitted curves created by Prism 9.2.0 (GraphPad Software). (d) Blood clearance profile of DiR-LNP in rats based on LNP-associated fluorescence obtained at 5 minutes, 30 minutes and 1 hour, with fitted curves created by Prism 9.2.0 (GraphPad Software). As the earliest three time points presented in **d** would become invisible if combined with 5 later time points, blood clearance profile of DiR-LNP based on all 8 time points was presented as two parts (**c** and **d**). Data in **c** and **d** were presented as “mean ± standard deviation” (n = 3) for each group, with differences between two injections analyzed using the multiple unpaired *t* test, with *P* values adjusted for FDR (false discovery rate). *, *P* < 0.05; **, *P* < 0.01; ***, *P* < 0.001; ****, *P* < 0.0001.

## Discussion

PEG-related immune responses induced by clinically relevant LNP may directly affect the clinical outcome of LNP-delivered drugs^11^. However, it is practically difficult to clarify these issues simply with clinical investigations. One major reason is the significant variability of pre-existing PEG-specific antibodies, making it extremely hard to identify anti-PEG antibodies specifically induced by LNP (**Supplementary Discussion**). Another concern is additional exposure to PEG derivatives other than LNP possibly existed during clinical observation period. In agreement with this, Ju *et al* reported that unvaccinated control donors had increased level of anti-PEG IgG and/or anti-PEG IgM in their clinical study^13^. Together with other above-mentioned influence factors existed in clinical studies such as small population size and varied mPEG_2000_ exposure amount, these limitations have led to inconsistent data regarding the characteristic of initial and repeated exposure to LNP-delivered drugs in inducing anti-PEG antibodies (**Supplementary Discussion**). Motivated by these challenges, we initiated the first animal study using a model LNP with the largest number of recipients all over the world, and delicately simulated clinical practice for both schedule and doses. Anti-PEG antibody was undetectable in all experimental groups before initial LNP injection, demonstrating a “clean” background of our model system. Encouragingly, through designing a series of time points and three doses (1:38:262) correlated with the amount of mPEG_2000_ contained in approved LNP-delivered drugs, we demonstrated for the first time that induction of anti-PEG IgM and IgG were both time- and dose-dependent, which are valuable for LNP-based research and development.

In spite of the significant difference between mRNA vaccines from traditional vaccines, no pharmacokinetic studies have been conducted for either approved mRNA vaccines or their LNP carriers^25,26^. Herein, we conducted the first pharmacokinetic study related with LNP-delivered vaccines or their LNP carriers. Our data suggest that previous LNP injection may accelerate the blood clearance of subsequently administered LNP, which has raised an important and clinically relevant issue that would warrant further investigation. It is noteworthy that although Alnylam Pharmaceuticals *Inc.* reported the absence of ABC phenomenon after repeated injection of Onpattro^®^, all patients in their study received corticosteroid premedication prior to each Onpattro^®^ injection to reduce the risk of infusion-related reactions^12^. However, corticosteroid is generally considered as an immunosuppressive drug and may repress PEG-associated immunological effects. To the best of our knowledge, previously there is no report investigating whether intramuscularly injected PEGylated therapeutics could induce accelerated blood clearance. Our data would thereby provide valuable information for revealing the pharmacokinetic characteristics of intramuscularly administered PEGylated therapeutics (**Supplementary Discussion**). Together with further in-depth studies using larger sample size, our findings may promote the optimization of premarket requirements and clinical practice for LNP-delivered biomedical products, as well as PEGylated products administered intramuscularly (**Supplementary Discussion**).

Similar to PEGylated liposome, LNP belongs to thymus-independent antigens (TI-Ag) as it contains no proteinatious composition^5,8^. There is a traditional perception that TI-Ag could not induce isotype switch from IgM to long-lasting IgG, resulting in IgM production only after TI-Ag administration, with no or very low level of IgG^8^. Moreover, different from thymus-dependent antigens (TD-Ag), TI-Ag generally could not induce B cell memory^27^. That is, an amplified, accelerated and affinity-matured antibody production could be observed after successive exposure to TD-Ag but not TI-Ag^27^. Consistent with these theories, even six repeated injections of PEGylated liposome neither enhanced anti-PEG IgM production nor effectively induced anti-PEG IgG in mice^28^. Unexpectedly, we discovered that LNP not only induced isotype switch and production of anti-PEG IgG, but caused B cell memory, leading to rapid enhancement and longer lasting time of both anti-PEG IgM and IgG upon repeated injections (**Supplementary** Fig. 8). To our best knowledge, there is no previous report on either inducing B cell memory or isotype switching from IgM to IgG by any PEGylated TI-Ag (**Supplementary Discussion**). These findings refine our understandings on PEGylated LNP, and possible other PEG derivatives.

## Methods

### Materials

Cholesterol (Cat. No. 57-88-5) and DSPC (Cat. No. 816-94-4) were purchased from Lipoid GMBH (Ludwigshafen, Germany). ALC-0315 (Cat. No. 06040008600) and ALC-0159 (Cat. No. 06020112302) were acquired from SINOPEG (Xiamen, China). Ferric chloride hexahydrate (Cat. No. 701122), ammonium thiocyanate (Cat. No. 221988) and NH_2_-PEG_10000_-NH_2_ (Cat. No. 8218815000) were obtained from Sigma-Aldrich (St. Louis, MO, USA). 3-[(3-cholamidopropyl)dimethylammonio]-1-propanesulfonate (CHAPS, Cat. No. ST1145), 3,3′,5,5′-tetramethylbenzidine dihydrochloride hydrate (TMB 2HCl, Cat. No. ST1708) and nonfat powdered milk (Cat. No. P0216) were purchased from Beyotime Biotechnology (Shanghai, China). 1,1-dioctadecyl-3,3,3,3-tetramethylindotricarbocyaine iodide (DiR, Cat. No. 100068-60-8) was purchased from Shanghai Maokang Biotechnology Co., Ltd. (Shanghai, China). Maxisorp 96-well microplates (Cat. No. 44-2404-21) were acquired from Nalge-Nunc International (Rochester, NY, USA). D-Luciferin (Cat. No. 88293) were purchased from Thermo Fisher Scientific (Waltham, MA, USA). Firefly luciferase mRNA (Cat. No. L-7702-1000) was obtained from Trilink Biotechnologies (San Diego, CA, USA). Rat anti-PEG IgM (Cat. No. rAGP6-PABM-A; Clone No. rAGP6) and rat anti-PEG IgG (Cat. No. r33G-PABG-A; Clone No. r33G) were acquired from Academia Sinica (Taipei, China). Peroxidase-conjugated affinipure rabbit anti-rat IgM µ-chain specific (Cat. No. 312-035-020; Clone No. 312-035-020) and peroxidase-conjugated affinipure donkey anti-rat IgG (H+L) (Cat. No. 712-035-150; Clone No. 712-035-150) were obtained from Jackson ImmunoResearch Laboratories Inc (West Grove, PA, USA).

### Preparation of LNP, DiR-LNP and DiR-LU@LNP

LNP, DiR-LNP and DiR-LU@LNP were formulated according to a previously reported protocol^29^. First, the ethanol phase was prepared by dissolving ALC-0315, DSPC, cholesterol and ALC-0159 at a molar ratio of 46.3: 9.4: 42.7: 1.6. Specifically, DiR was added into the ethanol phase at 0.4% mol for preparation of DiR-LNP and DiR-LU@LNP. Regarding the aqueous phase, it was prepared using 20 mM citrate buffer (pH4.0) for LNP and DiR-LNP formulations, with additional firefly luciferase mRNA added for DiR-LU@LNP formulation. Subsequently, the ethanol phase was mixed with the aqueous phase at a flow rate ratio of 1: 3 (ethanol: aqueous) through a microfluidic mixer (Precision Nanosystems Inc., Canada). Afterwards, the obtained nanoparticle solutions were dialyzed against 10 × volume of PBS (pH7.4) through a tangential-flow filtration (TFF) membrane with 100 kD molecular weight cut-off (Sartorius Stedim Biotech, Germany) for at least 18 hours. Finally, nanoparticle solutions were concentrated using Amicon ultra-centrifugal filters (EMD Millipore, Billerica, MA, USA), passed through a 0.22 µm filter and stored at 2∼8 ℃ until use.

### Characterization of LNP, DiR-LNP and DiR-LU@LNP

LNP, DiR-LNP and DiR-LU@LNP were examined for their hydrodynamic size (Z-average), polydispersity index (PDI) and zeta potential with DLS (Zetasizer Nano ZS, Malvern Instruments Ltd, Malvern, UK) equipped with a solid state HeNe laser (λ = 633 nm) at a scattering angle of 173°. Nanoparticles were either added into PBS (pH7.4) for Z-average and PDI measurements, or added into ultrapure water for determination of zeta potential. Three independent experiments were conducted, with each type of LNP examined at 25 ℃ for 10 seconds (pre-equilibration for 2 minutes) and repeated at least 10 times in disposable cuvettes (for Z-average and PDI) or zeta cuvettes (for zeta potential). The obtained data were presented as “mean ± standard deviation”. To further assess their stability in serum (simulating in vivo environment in this study), LNP, DiR-LNP and DiR-LU@LNP were diluted to 1:100 with PBS containing 10% rat serum and then incubated at 37 ℃ for 24 hours. Subsequently, 1 mL of diluted LNP, DiR-LNP and DiR-LU@LNP were respectively collected at designated time points (1 hour, 6 hours, 12 and 24 hours post-incubation), followed by characterization of Z-average and PDI with DLS. Three independent experiments were conducted, with each type of LNP examined at 37 ℃ for 10 seconds (pre-equilibration for 2 minutes) and repeated at least 10 times in disposable cuvettes. The obtained data were presented as “mean ± standard deviation”. Furthermore, the morphological characteristics of LNP, DiR-LNP and DiR-LU@LNP were observed with Cryo-TEM. In brief, 3 μL of each LNP sample was deposited onto a holey carbon grid that was glow-discharged (Quantifoil R1.2/1.3) and vitrificated using a Vitrobot Mark IV System (FEI/Thermo Scientific, Waltham, MA, USA). Cryo-TEM imaging was performed on a Talos F200C device (FEI/Thermo Scientific, Waltham, MA, USA) equipped with a 4k × 4k Ceta camera at 200 kV accelerating voltage in the Center of Cryo-Electron Microscopy, Zhejiang University.

In addition, the phospholipid (DSPC) concentrations of LNP, DiR-LNP and DiR-LU@LNP solutions were quantified via Steward’s assay for further calculation of LNP doses^30^. Briefly, ammonium ferrothiocynate was prepared by dissolving 27.03 mg ferric chloride hexahydrate and 30.4 mg ammonium thiocyanate in 1 mL of distilled water. 10 μL of the lipid sample was added to 990 μL of chloroform, followed by addition of 1 mL of ammonium ferrothiocynate. The obtained mixture was vortexed for 60 seconds and then centrifuged at 300 × g for 15 minutes at room temperature. The bottom chloroform layer was transferred to a glass cuvette and the absorbance was measured at 470 nm using a Unicam UV500 Spectrophotometer (Thermo electron corporation, USA). Standard curves for DSPC lipid were obtained and used for calculation of the phospholipid concentrations of LNP, DiR-LNP and DiR-LU@LNP solutions. Eventually, the various doses of LNP tested in the animal experiments were calculated based on the phospholipid (DSPC) exposure amount per dose of related drug (see below for details).

### Determination of LNP dosing protocols

According to the official drug information and clinical protocols, the mPEG_2000_ exposure amount for each injection of Comirnaty^®^ in adults is 0.0406 mg^18^, while that for Onpattro^®^ is 10.6434 mg^17^ (262 folds of that of Comirnaty^®^). Different from Comirnaty^®^ and Onpattro^®^, the detailed LNP composition of Spikevax^®^ including the molar lipid ratios is not included in the official drug information published in 2022^19^. With a postulation that PEG_2000_-DMG is the only lipid contained in LNP of Spikevax^®^, we estimated that the possible “maximum” mPEG_2000_ exposure for each injection would be 1.542 mg (37.98 folds of that of Comirnaty^®^) referred to the clinical protocols of Spikevax^®19^(**Supplementary Methods; Supplementary Table 1**).

Based on the above calculation and estimation, three mPEG_2000_ dosages including 0.0406 mg/dose (low dose), 1.542 mg/dose (middle dose) and 10.6434 mg/dose (high dose) were obtained. These dosages not only have an appropriate gradient ratio of 1:38:262, but cover the broad range of PEG exposure amount upon each injection of approved LNP-delivered drugs.

Next, we calculated the phospholipid (DSPC) contained in LNP of Comirnaty^®^, which is 0.09 mg for each injection in adults^18^. According to the animal-human dose exchange algorithm: animal equivalent dose = human dose × *K*_m_ ratio (6.2 for rat)^31^, clinically relevant LNP doses for rats were as follows: low dose (L-LNP), 0.009 mg phospolipid/kg (0.09 mg/60 kg × 6.2); middle dose (M-LNP), 0.342 mg phospholipids/kg (0.009 × 38); high dose (H-LNP), 2.358 mg phospholipids/kg (0.009 × 262).

The clinical protocols of Comirnaty^®^ were essentially simulated in this study. That is, LNP was administrated through intramuscular injection for two separate injections, with a 21-day interval (same as routine Comirnaty^®^ vaccination).

### Animals

10-12-week-old female Wistar rats were purchased from Hangzhou Medical College (Hangzhou, China), and maintained in the Laboratory Animal Center of Zhejiang University under controlled environmental conditions at constant temperature, humidity, and a 12-hour dark/light cycle. Rats were given ad libitum access to a standard rat chow and water, and were acclimated for at least 7 days. All animal experiments were approved by the Laboratory Animal Welfare and Ethnics Committee of Zhejiang University and carried out in accordance with the guidelines of the committee (approval No. ZJU20210071).

### Administration of LNP simulating clinical protocols and collection of serum samples for ELISA

Wistar rats were randomly divided into a Control group (n = 8) and three LNP-treated groups (n = 15). At Day 0, LNP-treated groups were intramuscularly injected with 0.009 mg phospholipids/kg LNP (L-LNP group), 0.342 mg phospholipids/kg LNP (M-LNP group) and 2.358 mg phospholipids/kg LNP (H-LNP group), respectively, while the Control group only received PBS. At Day 21, rats in each experimental group received same treatment as the initial injection. Peripheral blood samples of each rat were collected successively via the retro-orbital venous plexus at Day 0, 3, 5, 7, 14, 21, 24, 26, 28, 35, 42 and 49. All blood samples were centrifuged at 2000 × g for 15 minutes at 4 °C, and the serums were immediately stored at -80 °C for further quantification of anti-PEG antibody. At the experimental endpoint, the animals were humanely euthanized through intraperitoneal administration of 150mg/kg of sodium pentobarbital salt (Cat. No. BCP07810, Biochempartner, Shanghai, China)^32^.

### Quantification of anti-PEG IgM and anti-PEG IgG antibodies with ELISA

Maxisorp^®^ 96-well microplates were coated with 5 μg/well NH_2_-PEG_10000_-NH_2_ in 100 µL of PBS overnight at 4 ℃. Subsequently, plates were gently washed with 350 μL of washing buffer (0.05% (w/v) CHAPS in DPBS) for three times, followed by incubation with blocking buffer (5% (w/v) skim milk powder in DPBS, 200 μL/well) at room temperature for 1.5 hours. Afterwards, plates were rinsed with washing buffer for three times again. Then 100 μL of rat serum samples diluted 1: 150 with dilution buffer (2% (w/v) skim milk powder in DPBS), together with seven serial dilutions of rat anti-PEG IgM standards (1.37, 4.12, 12.35, 37.04, 111.11, 333.33 and 1000.00 ng/mL) or rat anti-PEG IgG standards (0.05, 0.15, 0.46, 1.37, 4.12, 12.35 and 37.04 ng/mL), were added into anti-PEG IgM or anti-PEG IgG detection plates in duplicate and further incubated for 1 hour at room temperature. After five successive washes, 50 µL of diluted peroxidase-conjugated affinipure rabbit anti-rat IgM µ-chain specific and peroxidase-conjugated affinipure donkey anti-rat IgG (H+L) antibodies were respectively added at 0.08 μg/mL to the corresponding plates and incubated for 1 hour at room temperature. Again, unbounded antibodies were removed by five washes, followed by incubation with 100 µL of TMB for 30 minutes at room temperature. Finally, HRP-TMB reaction was stopped with 100 μL of 2 N H_2_SO_4_, and the absorbance was measured at 450 nm with a microplate reader (Thermo Fisher Scientific, Waltham, MA, USA), using 570 nm as a reference wavelength. Anti-PEG IgM and anti-PEG IgG standard curves in each batch of ELISA were constructed by plotting the average corrected absorbance values (OD_450 nm_-OD_570 nm_) and corresponding antibody concentrations with Four Parameter Logistic (4PL) curve fit using Origin 2021 software (OriginLab Corporation, Northampton, Massachusetts, USA). The goodness of fit of each standard curve was measured by the coefficient of determination (*R*^2^)^33^. Concentrations of anti-PEG IgG and IgM antibodies in serum samples were calculated based on corresponding standard curves. In addition, intra-assay precision was evaluated by calculating the Coefficient of Variation (CV% = (Standard deviation/Mean) × 100%) for all detectable standards and samples in all batches of ELISA^23,24^. Inter-assay precision was determined by calculating the Coefficient of Variation for serially diluted anti-PEG antibody standards among all batches of ELISA^23,24^. The acceptance criteria for mean intra-assay and inter-assay Coefficient of Variation (CV%) of ELISA are < 20% and < 25%, respectively^23,24^.

### Biodistribution of PEGylated LNP in major organs of Wistar rats

Wistar rats were randomly divided into a Control group and three DiR-LU@LNP-treated groups (n = 6). At Day 0, LNP-treated groups were intramuscularly injected with 0.009 mg phospholipids/kg DiR-LU@LNP (L-LNP group), 0.342 mg phospholipids/kg DiR-LU@LNP (M-LNP group) and 2.358 mg phospholipids/kg DiR-LU@LNP (H-LNP group), respectively, while the Control group only received PBS. At Day 21, rats in each experimental group received same treatment as the initial injection. Six hours after the first and second injections, three rats in each group were administered intraperitoneally with D-luciferin at a dose of 150 mg/kg. Rats were sacrificed by cervical dislocation 15 minutes after D-luciferin administration and immediately dissected for collection of several primary organs, including heart, liver, spleen, lung, kidneys, draining lymph node and muscle at the injection site. Whole-organ/tissue imaginings for DiR fluorescence (Excitation/Emission: 748 nm/780 nm) and firefly luciferase bioluminescence were performed with IVIS Spectrum imaging system and analyzed with Living Image software (Caliper Life Sciences, Waltham, Massachusetts, USA).

### Blood clearance of PEGylated LNP in Wistar rats

Wistar rats were randomly divided into a Control group and three DiR-LNP-treated groups (n = 3). At Day 0, LNP-treated groups were intramuscularly injected with 0.009 mg phospholipids/kg DiR-LNP (L-LNP group), 0.342 mg phospholipids/kg DiR-LNP (M-LNP group) and 2.358 mg phospholipids/kg DiR-LNP (H-LNP group), respectively, while the Control group only received PBS. At Day 21, rats in each experimental group received same treatment as the initial injection. Peripheral blood samples were respectively collected from the retro-orbital venous plexus at 5 minutes, 30 minutes, 1 hour, 3 hours, 6 hours, 10 hours, 24 and 48 hours after the first and second injections. Then blood samples were centrifuged at 2000 × g at 4 °C for 15 minutes, and serum samples were isolated and immediately stored in dark at -80 °C. DiR fluorescence associated with LNP in serum samples was detected by fluorescent spectroscopy on a Spectramax ID5 (Molecular Devices, San Jose, California, USA) at excitation/emission wavelengths of 748/780 nm. At the experimental endpoint, the animals were humanely euthanized through intraperitoneal administration of 150mg/kg of sodium pentobarbital salt (Cat. No. BCP07810, Biochempartner, Shanghai, China)^32^.

### Data presentation and statistical analysis

All data were presented as “mean ± standard deviation”. Concentrations (ng/mL) of anti-PEG IgM and anti-PEG IgG were analyzed after log_10_ transformation, and their differences among various groups at each time point were analyzed with Mann-Whitney U test using R 4.0.5 (R Software, Boston, MA, USA), with *P* values adjusted with FDR (false discovery rate) method. Changing curves of average level of anti-PEG antibody over time for various doses were fitted by the R package called “ggalt”. Profile analysis was performed to examine whether the overall trends of changing curves of average level of anti-PEG antibody over time between every two groups were equal. The analysis included two parts: parallel test and coincidence test. Only when the two changing curves of average level of anti-PEG antibody met both parallel and coincidence test (*P* > 0.05), the overall trend of two changing curves of average anti-PEG antibody level was considered as no statistical difference. According to factorial design (group × time) and repeated measures of antibody level, linear mixed models (LMM) were conducted to compare the change rates and average levels of anti-PEG antibody across groups, with all time points included. Several variables, including group (indicating mean differences in the average levels of anti-PEG antibody), time, time^2^, number of injections, and interaction term of group and time (indicating mean differences in the change rates of anti-PEG antibody) as fixed effect and subject as random effect were considered in LMM.

In addition, ▴Anti-PEG IgM (Log_10_ CONC) was defined as Anti-PEG IgM (Log_10_ CONC_2nd injection_) (log_10_-transformed concentration of anti-PEG IgM induced during the second injection cycle) subtracting corresponding Anti-PEG IgM (Log_10_ CONC_1st injection_) (log_10_-transformed concentrations of anti-PEG IgM induced during the first injection cycle). Similarly, ▴Anti-PEG IgG (Log_10_ CONC) was calculated by subtracting Anti-PEG IgG (Log_10_ CONC_1st injection_) (log_10_-transformed concentrations of anti-PEG IgG induced during the first injection cycle) from the corresponding Anti-PEG IgG (Log_10_ CONC_2nd injection_) (log_10_-transformed concentration of anti-PEG IgG induced during the second injection cycle). Differences in ▴Anti-PEG IgM (Log_10_ CONC) or ▲Anti-PEG IgG (Log_10_ CONC) among various groups at each time point were analyzed with Mann-Whitney U test using R 4.0.5, with *P* values adjusted for FDR (false discovery rate). Changing curves of average level of ▲Anti-PEG IgM (Log_10_ CONC) or ▲Anti-PEG IgG (Log_10_ CONC) over time for various doses were fitted by the R package called “ggalt”. Profile analysis was performed to examine whether the overall trends of changing curves of average level of ▲Anti-PEG IgM (Log_10_ CONC) or ▲Anti-PEG IgG (Log_10_ CONC) over time between every two groups were equal. The analysis included two parts: parallel test and coincidence test. Only when the two changing curves of average level of ▲Anti-PEG IgM (Log_10_ CONC) or ▲Anti-PEG IgG (Log_10_ CONC) met both parallel and coincidence test, the overall trend of the two changing curves of average level of ▲Anti-PEG IgM (Log_10_ CONC) or ▲Anti-PEG IgG (Log_10_ CONC) was considered as no difference. According to factorial design (group × time) and repeated measures of antibody level, LMM were conducted to compare the change rates and average levels of ▲Anti-PEG IgM (Log_10_ CONC) or ▲Anti-PEG IgG (Log_10_ CONC) across groups, with all time points included. Several variables, including group (indicating mean differences in the average levels of ▲Anti-PEG IgM (Log_10_ CONC) or ▲Anti-PEG IgG (Log_10_ CONC)), time, time^2^, and interaction term of group and time (indicating mean differences in the change rates of ▲Anti-PEG IgM (Log_10_ CONC) or ▴Anti-PEG IgG (Log_10_ CONC) levels) as fixed effect and subject as random effect were considered in LMM. After performing the Shapiro-Wilk test to check for normality and the F test to check for variance homogeneity, data obtained in the biodistribution and blood clearance study were analyzed using multiple unpaired *t* tests with correction for multiple comparisons using Prism 9.2.0 (GraphPad Software, San Diego, USA). *P* < 0.05 was considered statistically significant.

## Data availability

All data supporting the findings of this study are available within the paper and Supplementary Information. The associated raw data are available from the corresponding author on reasonable request.

## Acknowledgments

The authors would like to gratefully acknowledge Shenghai Chang and Lingyun Wu in the Center of Cryo-Electron Microscopy (CCEM), Zhejiang University for their technical assistance on Cryo-EM. The authors would like to gratefully acknowledge Yanhong Chen in the Laboratory Animal Center, Zhejiang University for her technical assistance on animal experiments. The authors acknowledge support from the National Natural Science Foundation of China (21722405 and 22075243 to Dr. Meihua Sui), Zhejiang Provincial Natural Science Foundation of China (LZ23E030003 to Dr. Meihua Sui) and Startup Foundation for Hundred-Talent Program of Zhejiang University (to Dr. Meihua Sui).

## Author contributions

M.S., the principal investigator of the major supporting grants, and H.W. conceived the research project and defined the goals of the present study. M.S., H.W. and Y.W. designed the experiments. H.W. and Y.W. performed most experiments. M.S., H.W., Y.W., C.Y., X.X. and Y.H. discussed this project and analyzed the data. X.X. provided technical/platform supports. W.Z., H.L., Y.Z., G.L. and J.S. helped with the animal experiments. H.W., Y.W., C.Y. and Y.H. drafted the manuscript. M.S., H.W., Y.W., C.Y. and Y.H. revise and edited the manuscript. These authors contributed equally: H.W., Y.W. and C.Y.

## References

1. Mendes, B.B. et al. Nanodelivery of nucleic acids. Nat. Rev. Methods Primers 2, 24 (2022).

2. Curreri, A., Sankholkar, D., Mitragotri, S. & Zhao, Z. RNA therapeutics in the clinic. Bioeng. Transl. Med. e10374 (2022).

3. Suk, J.S., Xu, Q.G., Kim, N., Hanes, J. & Ensign, L.M. PEGylation as a strategy for improving nanoparticle-based drug and gene delivery. Adv. Drug Deliver. Rev. 99, 28–51 (2016).

4. Khurana, A. et al. Role of nanotechnology behind the success of mRNA vaccines for COVID-19. Nano Today 38, 101142 (2021).

5. Chen, B.M., Cheng, T.L. & Roffler, S.R. Polyethylene glycol immunogenicity: theoretical, clinical, and practical aspects of anti-polyethylene glycol antibodies. ACS Nano 15, 14022–14048 (2021).

6. Ibrahim, M. et al. Polyethylene glycol (PEG): the nature, immunogenicity, and role in the hypersensitivity of PEGylated products. J. Control. Release 351, 215–230 (2022).

7. Yang, Q. & Lai, S.K. Anti-PEG immunity: emergence, characteristics, and unaddressed questions. Wiley Interdiscip. Rev. Nanomed. Nanobiotechnol. 7, 655–677 (2015).

8. Shi, D. et al. To PEGylate or not to PEGylate: immunological properties of nanomedicine’s most popular component, polyethylene glycol and its alternatives. Adv. Drug Deliver. Rev. 180, 114079 (2022).

9. Chen, B.M. et al. Measurement of pre-existing IgG and IgM antibodies against polyethylene glycol in healthy individuals. Anal. Chem. 88, 10661–10666 (2016).

10. Yang, Q. et al. Analysis of pre-existing IgG and IgM antibodies against polyethylene glycol (PEG) in the general population. Anal. Chem. 88, 11804–11812 (2016).

11. Ju, Y. et al. Impact of anti-PEG antibodies induced by SARS-CoV-2 mRNA vaccines. Nat. Rev. Immunol. 23, 1–2 (2022).

12. Zhang, X.P. et al. Patisiran pharmacokinetics, pharmacodynamics, and exposure-response analyses in the phase 3 APOLLO trial in patients with hereditary transthyretin-mediated (hATTR) amyloidosis. J. Clin. Pharmacol. 60, 37–49 (2020).

13. Ju, Y. et al. Anti-PEG antibodies boosted in humans by SARS-CoV-2 lipid nanoparticle mRNA vaccine. ACS Nano 16, 11769–11780 (2022).

14. Guerrini, G. et al. Monitoring anti-PEG antibodies level upon repeated lipid nanoparticle-based COVID-19 vaccine administration. Int. J. Mol. Sci. 23, 8838 (2022).

15. Carreño, J.M. et al. mRNA-1273 but not BNT162b2 induces antibodies against polyethylene glycol (PEG) contained in mRNA-based vaccine formulations. Vaccine 40, 6114–6124 (2022).

16. Bavli, Y., et al. Anti-PEG antibodies before and after a first dose of Comirnaty (mRNA-LNP-based SARS-CoV-2 vaccine). J. Control. Release 354, 316–322 (2023).

17. ONPATTRO-Prescribing Information, https://www.alnylam.com/sites/default/files/pdfs/ONPATTRO-Prescribing-Information.pdf (2023).

18. Food and Drug Administration. Comirnaty Information-Summary basis for regulatory action, 8 November, 2021; https://www.fda.gov/media/151733/download

19. Food and Drug Administration. Spikevax Information-Summary basis for regulatory action, 30 January, 2022; https://www.fda.gov/media/155931/download

20. World Health Organization. WHO guidelines on non-clinical evaluation of vaccines, WHO Technical Report Series No. 927, Annex 1, 1 January 2005; https://cdn.who.int/media/docs/default-source/biologicals/vaccine-standardization/annex-1nonclinical.p31-63.pdf?sfvrsn=e87c28d8_3&download=true

21. Yang, K., Reker-Smit, C., Stuart, M.C.A. & Salvati, A. Effects of protein source on liposome uptake by cells: corona composition and impact of the excess free proteins. Adv. Healthc. Mater. 10, e2100370 (2021).

22. Naumenko, V.A. et al. Intravital imaging of liposome behavior upon repeated administration: a step towards the development of liposomal companion diagnostic for cancer nanotherapy. J. Control. Release 330, 244–256 (2021).

23. Food and Drug Administration. Guidance for industry: bioanalytical method validation. http://www.fda.gov/downloads/Drugs/GuidanceComplianceRegulatoryInformation/Guidances/ucm070107.pdf. (2018)

24. Findlay, J.W.A. et al. Validation of immunoassays for bioanalysis: a pharmaceutical industry perspective. J. Pharmaceut. Biomed. 21, 1249–1273 (2000).

25. European Medicines Agency. Spikevax (previously COVID-19 Vaccine Moderna) : EPAR - Public assessment report, 20 January 2021; https://www.ema.europa.eu/en/documents/assessment-report/spikevax-previously-covid-19-vaccine-moderna-epar-public-assessment-report_en.pdf

26. European Medicines Agency. Comirnaty : EPAR - Public assessment report, 23 December 2020; https://www.ema.europa.eu/en/documents/assessment-report/comirnaty-epar-public-assessment-report_en.pdf

27. Defrance, T., Taillardet, M. & Genestier, L. T cell-independent B cell memory. Curr. Opin. Immunol. 23, 330–336 (2011).

28. Ichihara, M. et al. Anti-PEG IgM response against PEGylated liposomes in mice and rats. Pharmaceutics 3, 1–11 (2010).

29. Zhang, N.N. et al. A thermostable mRNA vaccine against COVID-19. Cell 182, 1271–1283 (2020).

30. Stewart, J.C.M. Colorimetric determination of phospholipids with ammonium ferrothiocyanate. Anal. Biochem. 104, 10–14 (1980).

31. Nair, A., Morsy, M.A. & Jacob, S. Dose translation between laboratory animals and human in preclinical and clinical phases of drug development. Drug Develop. Res. 79, 373–382 (2018).

32. American Veterinary Medical Association. AVMA guidelines for the euthanasia of animals: 2020 Edition, 24 February 2020; https://www.avma.org/sites/default/files/2020-02/Guidelines-on-Euthanasia-2020.pdf

33. Draper, N. R., & Smith, H. Applied regression analysis (3rd ed.) (Wiley Press, New York, 1998).

