## Supplementary Materials for "Polyethylene glycol (PEG)-associated immune responses triggered by clinically relevant lipid nanoparticles in rats"

### Supplementary Discussion

#### Significant person-to-person and study-to-study variabilities in pre-existing anti-PEG antibodies

PEG is a versatile polymer commonly used as a surfactant, solvent and emulsifying agent in household chemicals, as an additive in foods, and as either an active composition or an inactive excipient in medicine[1]. Currently FDA has approved 33 PEGylated agents for a variety of clinical indications such as metabolic disease, immunological disease, degenerative disease, cancer and infectious diseases (https://www.drugs.com). Since anti-PEG IgM was first detected in rabbits immunized with PEGylated ovalbumin in 1983[2], an expanding body of evidence has revealed that some PEG derivatives could elicit PEG-specific antibodies[3-5]. Interestingly, some people who never received PEGylated drugs have pre-existing antibodies against PEG possibly due to environmental exposure[4,5]. For instance, an epidemiological study based on 1504 healthy Han Chinese donors residing in Taiwan area of China found that a total of 666 individuals (44.3%) had positive anti-PEG IgG or IgM, with 25.7%, 27.1%, and 8.4% of the total population having anti-PEG IgG only, anti-PEG IgM only, and both anti-PEG IgG and IgM, respectively[6]. This study also showed that PEG-specific antibodies were more common in females than in males (32.0% *vs* 22.2% for IgM and 28.3% *vs* 23.0% for IgG), and in young people (up to 60% for 20 years old) as compared to old people (20% for > 50 years old). Another epidemiological study based on 377 healthy human blood donors in USA found that anti-PEG antibodies were detectable in ~72% of individuals, with 18%, 25% and 30% of all samples having anti-PEG IgG only, anti-PEG IgM only, and both anti-PEG IgG and IgM, respectively[7].

Up to date there are five published studies that evaluated the induction of anti-PEG antibodies by approved LNP-delivered drugs, including three related with Comirnaty®, Spikevax® and mixed use of these two vaccines[8-11]. However, it is noteworthy that these limited available literature showed significant study-to-study variability in pre-existing anti-PEG antibody: Alnylam Pharmaceuticals *Inc.* reported that only two of 224 patients (0.89%) with hereditary transthyretin-mediated (hATTR) amyloidosis were positive for anti-PEG antibodies at baseline[8]; Ju *et al* from the University of Melbourne stated that anti-PEG IgG was commonly detectable (71%) before vaccination in Comirnaty® and Spikevax® cohorts[9]; Guerrini *et al* from Joint Research Centre in Italy described that anti-PEG IgG was positive before the first vaccine injection in their cohorts receiving two LNP-based COVID-19 vaccines, with a large person-to-person variability[10]. Carreño *et al* from Icahn School of Medicine at Mount Sinai in USA did not report the status of pre-existing anti-PEG antibodies in their very small population study (n = 10)[11]. Bavli *et al* from Hebrew University-Hadassah Medical School in Israel showed that anti-PEG IgG, IgM and IgE was detected in 29 (36.7%), 11 (13.9%) and 0 individuals, respectively, before vaccination with Comirnaty®[12].These significant variabilities in pre-existing anti-PEG antibodies would lead to unfavorable intervention when identifying and analyzing antibodies induced by PEGylated LNP.

#### Inconsistent previous results regarding the induction of anti-PEG antibodies by PEGylated LNP-delivered therapeutics

Across very limited population-based studies, no consistent results was obtained regarding any characteristic of initial and/or repeated injection of LNP-delivered drugs in inducing any type of antibodies against PEG:

Alnylam Pharmaceuticals *Inc.* reported that anti-PEG IgM and IgG were induced in 3.4% of subjects (5 out of 145 patients) who received Onpattro® in 2019[8]; Ju *et al* reported in 2022 that COVID-19 mRNA vaccines boosted the serum anti-PEG antibody levels in Australian recipients, with anti-PEG IgM boosted a mean of 2.64 folds and anti-PEG IgG boosted a mean of 1.78 folds following Comirnaty® vaccination (n = 55), as well as anti-PEG IgM boosted a mean of 68.5 folds and anti-PEG IgG boosted a mean of 13.1 folds following Spikevax® vaccination (n = 20)[9]; Guerrini *et al* from Joint Research Centre in Italy reported a significant increase in anti-PEG IgM level after the first injection of Comirnaty® and the third injection of Comirnaty® or Spikevax®, while no boosting effect was observed on anti-PEG IgG after injection with either vaccine in 2022[10]; Carreño *et al* reported different response on induction of PEG-specific antibodies with a very small size of recipients in USA received either Comirnaty® or Spikevax® vaccination (n = 10) in 2022[11]. Besides, the fold changes of both anti-PEG IgM and IgG induced by either mRNA vaccine had a very broad range. As stated by the authors, small population sizes, pre-existing antibodies, inevitable interference due to exposure to PEG-containing substances other than vaccines after immunization, as well as other potential influence factors, may affect the reliability of their data[9,11]. Bavli *et al* from Israel reported a significant increase in serum anti-PEG IgG three weeks after the first Comirnaty® administration, while no increase in anti-PEG IgM or IgE was detected (n = 79)[12].

**Additional interpretation of accelerated blood clearance induced by repeated injection of PEGylated LNP intramuscularly**

It is well known that intramuscular administration results in drug absorption and clearance significantly different from intravenous injection[13,14]. For instance, intravenously administered drugs immediately enter the blood circulation and reach the maximum blood concentration (Cmax). Therefore, accelerated blood clearance (ABC) phenomenon could be observed right after repeated intravenous injection of PEGylated drugs due to the instant "antigen-antibody" binding in the blood[15,16]. However, it takes a while for intramuscularly injected drugs to be absorbed from injection site into the blood to reach the Cmax[13,14]. It is thus understandable that accelerated blood clearance induced by H-LNP re-injection was observed at 30 minutes and 60 minutes after intramuscular reinjection, rather than at the earliest time point such as 5 minutes (**Fig. 4c**). On the other hand, after "neutralization" of circulating anti-PEG antibodies by newly injected LNP, or the remaining "antigen-antibody" binding is not abundant enough to significantly reduce LNP-associated fluorescence in circulation, the blood clearance will return to normal. Thenceforth LNP absorbed from intramuscular injection site into blood could gradually increase LNP-associated fluorescence. For instance, peak level of fluorescence reached at around 24 hours after repeated injection of H-LNP (**Fig. 4c**). Interestingly, ABC phenomenon arose again at 48 hours after repeated injection of H-LNP, which coincides with the correspondingly enhanced production of anti-PEG IgM and IgG antibodies at this time point (**Figs. 2-4**).

It is noteworthy that the levels of "pre-existing" anti-PEG antibodies are expected to be gradually increased with a higher number of repeated LNP injections. This may lead to occurrence of accelerated blood clearance even in L-LNP and M-LNP groups, as well as a more pronounced ABC phenomenon in the H-LNP group. Considering that Onpattro® needs to be continuously/repeatedly injected until the patient's condition is ideally controlled, and that both COVID-19 mRNA vaccines are used for booster immunization after routine two-injection vaccination, our findings may have broad clinical implications.

**Unexpected induction of B cell memory and isotype switching by PEGylated LNP**

Our model system has provided an opportunity to explore the mechanisms mediating the generation of anti-PEG antibodies induced by clinically relevant LNP. It is well known that non-protein antigens, such as lipids, polysaccharides, and naturally occurring non-proteinatious and synthetic polymers, can stimulate antibody response in the absence of T helper cell and is therefore called thymus-independent antigens or T cell-independent antigens (TI-Ag)[3,17]. In contrast, T-dependent antigens (TD-Ag) mainly include proteins/peptides that are taken up by the antigen-presenting cells and presented in the context with major histo-compatibility complex type 2 (MHC II) to the T helper lymphocytes[3,-17]. According to its chemical nature, LNP is similar to PEGylated liposome and belongs to TI-Ag. It is generally believed that TI-Ag could induce neither isotype switch from IgM to long-lasting IgG nor a typical recall antibody response, which is also called B cell memory characterized by an amplified, accelerated and affinity-matured antibody production after successive exposure to certain antigens such as TD-Ag[17-19]. After a thorough literature search, we found that although three types of TI-Ag, including B. hermsii (Borrelia hermsii, a relapsing fever bacterium), NP-Ficoll (4-hydroxy-3-nitrophenylacetyl-Ficoll, a model TI-Ag) and pneumococcal capsular PS3 (serotype 3 capsular polysaccharide), could induce B cell memory[20-22], previously there is no report on either inducing B cell memory or isotype switching by any PEG derivatives belonging to TI-Ag. It needs to be pointed out that no related conclusion could be drawn from the above-mentioned four clinical studies evaluating anti-PEG antibodies induced by LNP-delivered drugs, as the necessary statistical analysis on anti-PEG antibody production was not conducted in all these reports. Herein, our data showing induction of isotype switching from anti-PEG IgM to IgG, as well as B cell memory by repeated LNP injection, has revealed new immune properties of PEGylated LNP (**Supplementary Fig. 8**).

Considering the huge population exposed to clinically relevant LNP (total sales volume of Comirnaty® > 5,341,276,760 doses; total sales volume for Spikevax® > 3,229,743,423 doses; from WHO website (https://app.powerbi.com/view?r=eyJrIjoiMWNjNzZkNjctZTNiNy00YmMzLTkxZjQtNmJiZDM2MTYxNzEwIiwidCI6ImY2MTBjMGI3LWJkMjQtNGIzOS04MTBiLTNkYzI4MGFmYjU5MCIsImMiOjh9), and the rapid development of LNP-based therapeutics, further studies on PEG-associated immune responses triggered by LNP are warranted.

### Supplementary Methods

**Additional information for determination of clinically relevant $mPEG_{2000}$ and LNP dose gradients**

Complete LNP composition of Comirnaty® and Onpattro® can be respectively found in the following links: Food and Drug Administration. Comirnaty Information-Summary basis for regulatory action, 8 November, 2021, https://www.fda.gov/media/151733/download; https://www.alnylam.com/sites/default/files/pdfs/ONPATTRO-Prescribing-Information.pdf. However, although the LNP composition of mRNA-1273 used in a preclinical study was reported previously[23], this recipe has not been confirmed by the official drug instructions from FDA and

Moderna *Inc.* published later: Food and Drug Administration. Spikevax Information-Summary basis for regulatory action, 30 January, 2022, https://www.fda.gov/media/155931/download. As the detailed LNP formulation of Spikevax® has been kept confidential till now, alternatively two calculation or estimation methods through which an appropriate middle exposure dose of $mPEG_{2000}$ was determined (**Supplementary Table 1**). Eventually, clinically relevant $mPEG_{2000}$ and corresponding LNP dosages were determined, with an appropriate gradient ratio of 1:38:262 (see context).

**Supplementary Table 1. Determination of clinically relevant $mPEG_{2000}$ and LNP dose gradients***

| Approved therapeutics | Detailed LNP composition in official drug instructions | $mPEG_{2000}$ dose in adult | Relation to dose gradients | Equivalent LNP dose in rat |
|---|---|---|---|---|
| BNT162b2 /Comirnaty® | 0.43 mg/dose ALC-0315; 0.05 mg/dose ALC-0159; 0.09 mg/dose DSPC; 0.2 mg/dose Cholesterol | 0.0406 mg/dose based on official drug instructions | Precisely related to L-LNP | 0.009 mg phospholipid/kg |
| mRNA-1273 /Spikevax® | The only preclinical study published in 2020 introduced the molar lipid ratios (%) (ionizable cationic lipid: PEGylated lipid: DSPC: Cholesterol) of LNP are 50:1.5:10:38.5. | 0.093 mg/dose (2.3 folds of that of Comirnaty®) based on a LNP recipe described in a preclinical study with no further confirmation by official drug instructions | No relation | N/A |
| | Officially FDA and Moderna *Inc.* only described the total content of lipids (1.93 mg/dose) that make up LNP, while kept the detailed composition including the molar lipid ratios confidential till now. | 1.542 mg/dose (38 folds of that of Comirnaty®; possible "maximum" exposure) based on a postulation that $PEG_{2000}$-DMG is the only lipid contained in LNP | Related to M-LNP | 0.342 mg phospholipid/kg (0.009 × 38) |
| Patisiran /Onpattro® | 117 mg/dose DLin-MC3-DMA; 14.4 mg/dose PEG2000-C-DMG; 29.7 mg/dose DSPC; 55.8 mg/dose Cholesterol | 10.6434 mg/dose (262 folds of that of Comirnaty®) based on official drug instructions | Precisely related to H-LNP | 2.358 mg phospholipid/kg (0.009 × 262) |

***Animal-human dose exchange algorithm: animal equivalent dose=human dose × *K*m ratio (6.2 for rat)**

### Supplementary References


1. Ibrahim, M. *et al*. Polyethylene glycol (PEG): the nature, immunogenicity, and role in the hypersensitivity of PEGylated products. *J. Control. Release* **351**, 215-230 (2022).
2. Richter, A.W. & Akerblom, E. Antibodies against polyethylene-glycol produced in animals by immunization with monomethoxy polyethylene-glycol modified proteins. *Int. Arch. Aller. A. Imm.* **70**, 124-131 (1983).
3. Chen, B.M., Cheng, T.L. & Roffler, S.R. Polyethylene glycol immunogenicity: theoretical, clinical, and practical aspects of anti-polyethylene glycol antibodies. *ACS Nano* **15**, 14022-14048 (2021).
4. Haddad, H.F., Burke, J.A., Scott, E.A. & Ameer, G.A. Clinical relevance of pre-existing and treatment-induced anti-poly(ethylene glycol) antibodies. *Regen. Eng. Transl. Med.* **8**, 32-42 (2022).
5. Thi, T.T.H. *et al*. The importance of poly(ethylene glycol) alternatives for overcoming PEG immunogenicity in drug delivery and bioconjugation. *Polymers-Basel* **12**, 298 (2020).
6. Chen, B.M. *et al*. Measurement of pre-existing IgG and IgM antibodies against polyethylene glycol in healthy individuals. *Anal. Chem.* **88**, 10661-10666 (2016).
7. Yang, Q. *et al*. Analysis of pre-existing IgG and IgM antibodies against polyethylene glycol (PEG) in the general population. *Anal. Chem.* **88**, 11804-11812 (2016).
8. Zhang, X.P. *et al*. Patisiran pharmacokinetics, pharmacodynamics, and exposure-response analyses in the phase 3 APOLLO trial in patients with hereditary transthyretin-mediated (hATTR) amyloidosis. *J. Clin. Pharmacol.* **60**, 37-49 (2020).
9. Ju, Y. *et al*. Anti-PEG antibodies boosted in humans by SARS-CoV-2 lipid nanoparticle mRNA vaccine. *ACS Nano* 1**6**, 1769-11780 (2022).

10. Guerrini, G. *et al*. Monitoring anti-PEG antibodies level upon repeated lipid nanoparticle-based COVID-19 vaccine administration. *Int. J. Mol. Sci.* **23**, 8838 (2022).
11. Carreño, J.M. *et al*. mRNA-1273 but not BNT162b2 induces antibodies against polyethylene glycol (PEG) contained in mRNA-based vaccine formulations. *Vaccine* **40**, 6114-6124 (2022).
12. Bavli, Y. *et al*. Anti-PEG antibodies before and after a first dose of Comirnaty (mRNA-LNP-based SARS-CoV-2 vaccine). *J. Control. Release* **354**, 316-322 (2023).
13. León-Tamariz, F. *et al.* Biodistribution and pharmacokinetics of PEG-10kDa-cholecystokinin-10 in rats after different routes of administration. *Curr. Drug Deliv.* **7**, 137-143 (2010).
14. Choi, H.K. *et al.* In vitro and in vivo study of poly(ethylene glycol) conjugated ketoprofen to extend the duration of action. *Int. J. Pharm.* **341**, 50-57 (2007).
15. Ishida, T. *et al.* Accelerated blood clearance of PEGylated liposomes following preceding liposome injection: effects of lipid dose and PEG surface-density and chain length of the first-dose liposomes. *J. Control. Release* **105**, 305-317 (2005).
16. Liu, M. *et al.* A preliminary study of the innate immune memory of Kupffer cells induced by PEGylated nanoemulsions. *J. Control. Release* **343**, 657-671 (2022).
17. Shi, D. *et al*. To PEGylate or not to PEGylate: immunological properties of nanomedicine's most popular component, polyethylene glycol and its alternatives. *Adv. Drug Deliver. Rev.* **180**, 114079 (2022).
18. Kozma, G.T., Shimizu, T., Ishida, T. & Szebeni, J. Anti-PEG antibodies: properties, formation, testing and role in adverse immune reactions to PEGylated nano-biopharmaceuticals. *Adv. Drug Deliv. Rev.***154-155**, 163-175 (2020).
19. Defrance, T., Taillardet, M. & Genestier, L. T cell-independent B cell memory. *Curr. Opin. Immunol.* **23**, 330-336 (2011).
20. Alugupalli, K.R. *et al*. B1b lymphocytes confer T cell-independent long-lasting immunity. *Immunity* **21**, 379-390 (2004).
21. Obukhanych, T.V. & Nussenzweig, M.C. T-independent type II immune responses generate memory B cells. *J. Exp. Med.* **203**, 305-310 (2006).
22. Taillardet, M. *et al*. The thymus-independent immunity conferred by a pneumococcal polysaccharide is mediated by long-lived plasma cells. *Blood* **114**, 4432-4440 (2009).
23. Corbett, K.S. *et al*. SARS-CoV-2 mRNA vaccine design enabled by prototype pathogen preparedness. *Nature* **586**, 567-571 (2020).

**Supplementary Table 2. Characterization of LNP, DiR-LNP and DiR-LU@LNP**

| Formulation | Z-average (nm) | PDI | Zeta potential (mV) |
|---|---|---|---|
| LNP | 110.400 ± 3.466 | 0.203 ± 0.012 | 16.733 ± 0.451 |
| DiR-LNP | 113.067 ± 2.139 | 0.183 ± 0.013 | 7.257 ± 0.168 |
| DiR-LU@LNP | 101.367 ± 2.593 | 0.197 ± 0.015 | -5.943 ± 0.129 |

Data were presented as “mean ± standard deviation” of three independent experiments.

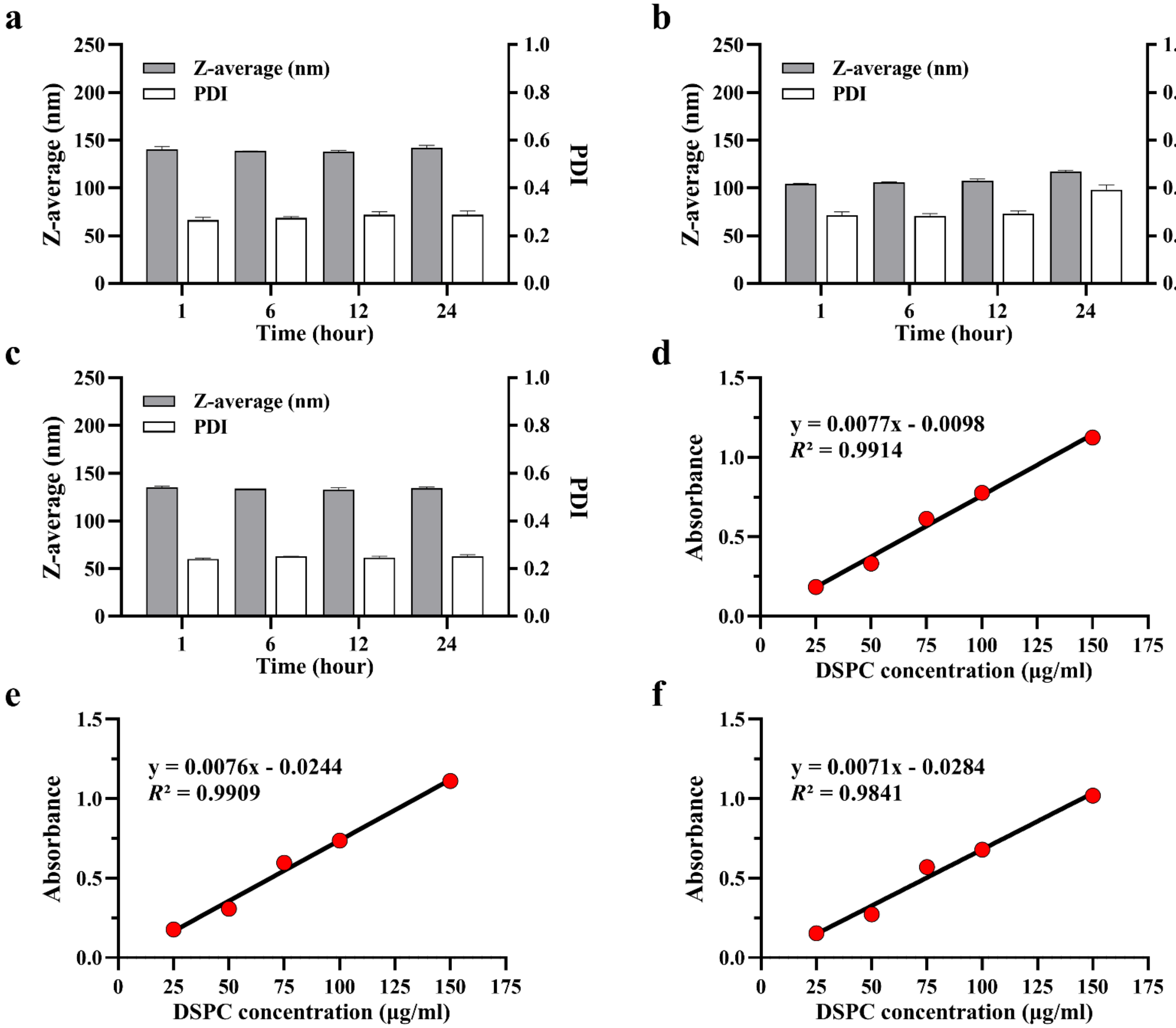


**Supplementary Fig. 1. Stability of LNP, DiR-LNP and DiR-LU@LNP in serum and standard curves for phospholipid (DSPC).** (**a-c**) Stability of (a) LNP, (b) DiR-LNP and (c) DiR-LU@LNP in serum. LNP, DiR-LNP and DiR-LU@LNP were diluted to 1:100 with PBS containing 10% rat serum and incubated at 37°C for 24 h. Subsequently, 1 mL of diluted LNP, DiR-LNP and DiR-LU@LNP were respectively collected at designated time points (1 h, 6 h, 12 h and 24 h), followed by characterization of Z-average and PDI with dynamic light scattering. Z-average/PDI of three LNP formulations at four successive time points were as follows: LNP, 140.533 ± 2.768 nm/0.264 ± 0.012, 138.600 ± 0.100 nm/0.274 ± 0.005, 138.200 ± 0.954 nm/0.287 ± 0.013 and 141.867 ± 2.631 nm/0.287 ± 0.016; DiR-LNP, 104.300 ± 0.458 nm/0.285 ± 0.014, 105.733 ± 0.503 nm/0.282 ± 0.010, 107.267 ± 1.940 nm/0.291 ± 0.013 and 117.200 ± 1.277 nm/0.392 ± 0.020; DiR-LU@LNP, 135.067 ± 1.550 nm/0.240 ± 0.003, 133.867 ± 0.058 nm/0.251 ± 0.001, 132.667 ± 2.023 nm/0.246 ± 0.006 and 134.133 ± 1.222 nm/0.252 ± 0.006. (**d-f**) Standard curves for determining phospholipid (DSPC) concentration in (d) LNP, (e) DiR-LNP and (f) DiR-LU@LNP solutions. Correspondingly, following equations were respectively obtained, in which y represents absorbance measured at 470 nm and x represents phospholipid concentration: LNP, y = 0.0077x + 0.0098 ($R^2$ = 0.9914); DiR-LNP, y = 0.0076x + 0.0244 ($R^2$ = 0.9909); DiR-LU@LNP, y = 0.0071x + 0.0284 ($R^2$ = 0.9841). Data were presented as "mean ± standard deviation" of three independent experiments.

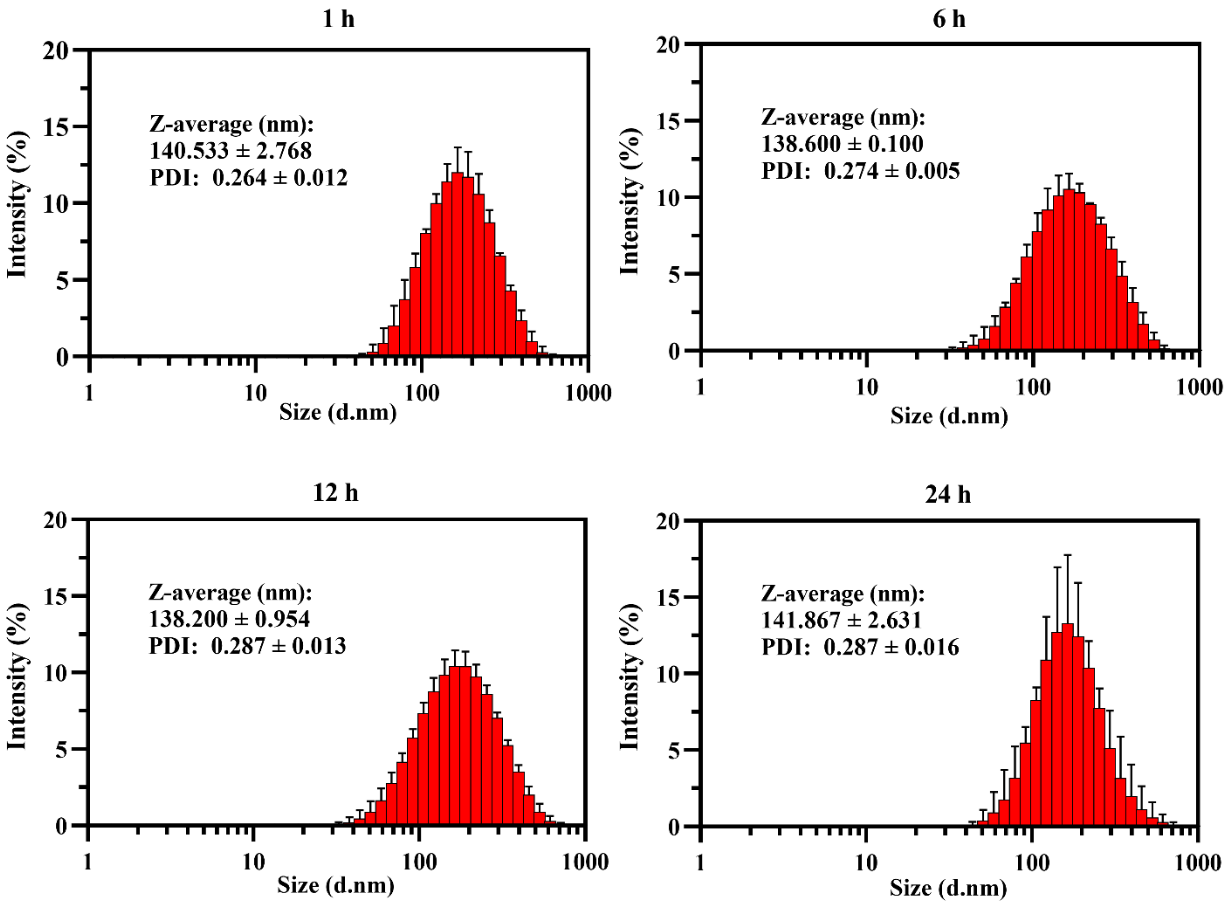


**Supplementary Fig. 2. Determination of LNP stability in serum.** LNP was diluted to 1:100 with PBS containing 10% rat serum and incubated at 37 °C for 24 h. Subsequently, 1 mL of diluted LNP was collected at designated time points (1 h, 6 h, 12 h and 24 h), followed by characterization of Z-average and PDI with dynamic light scattering. Data were presented as "mean ± standard deviation" of three independent experiments.

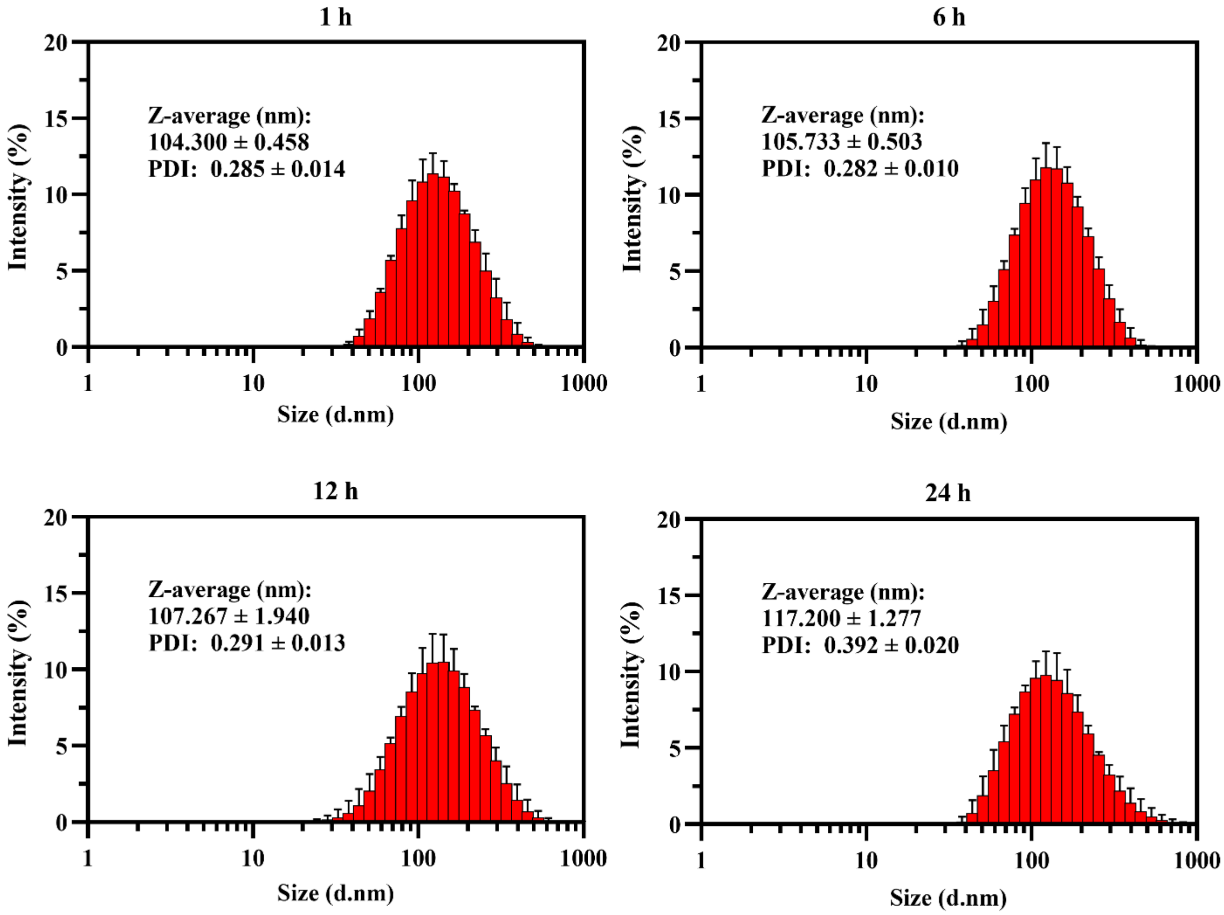


**Supplementary Fig. 3. Determination of DiR-LNP stability in serum.** DiR-LNP was diluted to 1:100 with PBS containing 10% rat serum and incubated at 37 ℃ for 24 h. Subsequently, 1 mL of diluted LNP was collected at designated time points (1 h, 6 h, 12 h and 24 h), followed by characterization of Z-average and PDI with dynamic light scattering. Data were presented as "mean ± standard deviation" of three independent experiments.

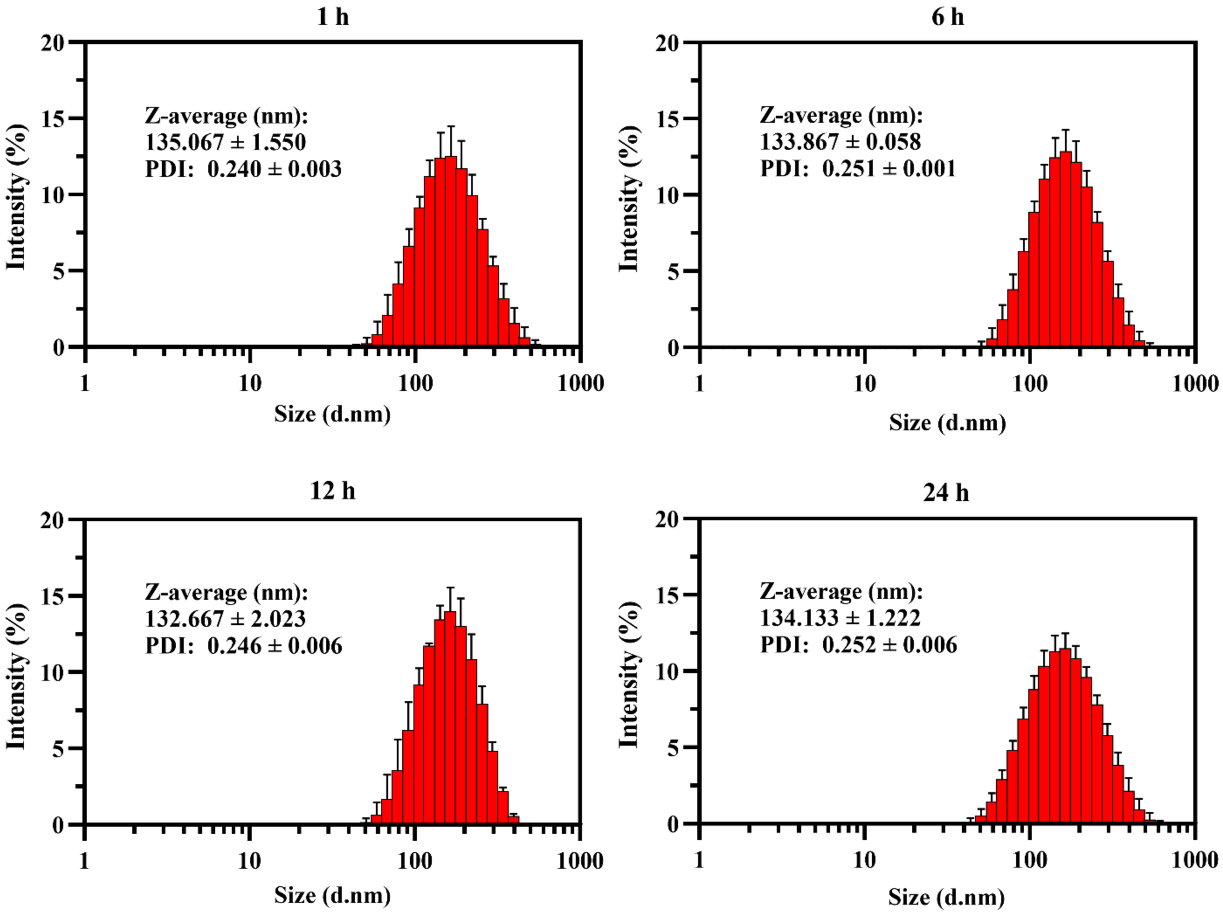


**Supplementary Fig. 4. Determination of DiR-LU@LNP stability in serum.** DiR-LU@LNP was diluted to 1:100 with PBS containing 10% rat serum and incubated at 37 °C for 24 h. Subsequently, 1 mL of diluted LNP was collected at designated time points (1 h, 6 h, 12 h and 24 h), followed by characterization of Z-average and PDI with dynamic light scattering. Data were presented as "mean ± standard deviation" of three independent experiments.

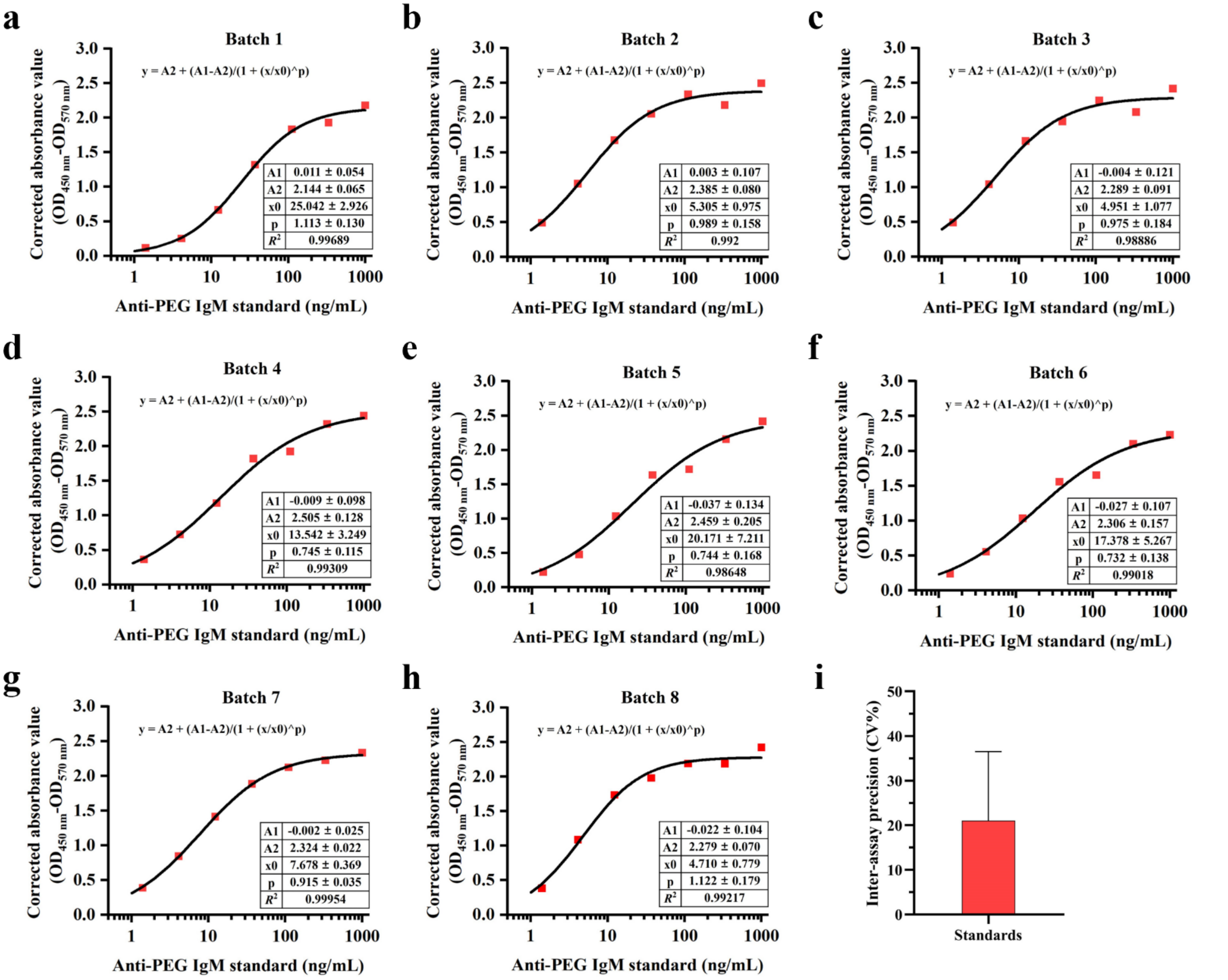


**Supplementary Fig. 5. Standard curves of ELISA for detecting anti-PEG IgM in rat serum samples (a-h) and inter-assay precision (CV%) of anti-PEG IgM standards (i).** Standard curves were constructed by plotting the average absorbance values ($OD_{450\ nm}$-$OD_{570\ nm}$) and corresponding antibody concentrations with Four Parameter Logistic (4PL) curve fit using Origin 2021 software. Serial dilutions of anti-PEG IgM standards (1.37, 4.12, 12.35, 37.04, 111.11, 333.33 and 1000.00 ng/mL) were included in each batch of ELISA for total eight independent batches. Inter-assay precision was determined by calculating the Coefficient of Variation (CV% = (Standard deviation/Mean) ×100%) for anti-PEG IgM standards among all eight batches of ELISA, which was 20.983 ± 15.511% as indicated in subfigure **i** (see **Methods** for acceptance criteria). In addition to the anti-PEG IgM standards run for each batch, 88 different rat serum samples were respectively tested in batch 1-3 and batch 5-7, and 54 different rat serum samples were respectively tested in batch 4 and 8. Data in **i** were presented as "mean ± standard deviation" (n = 7).

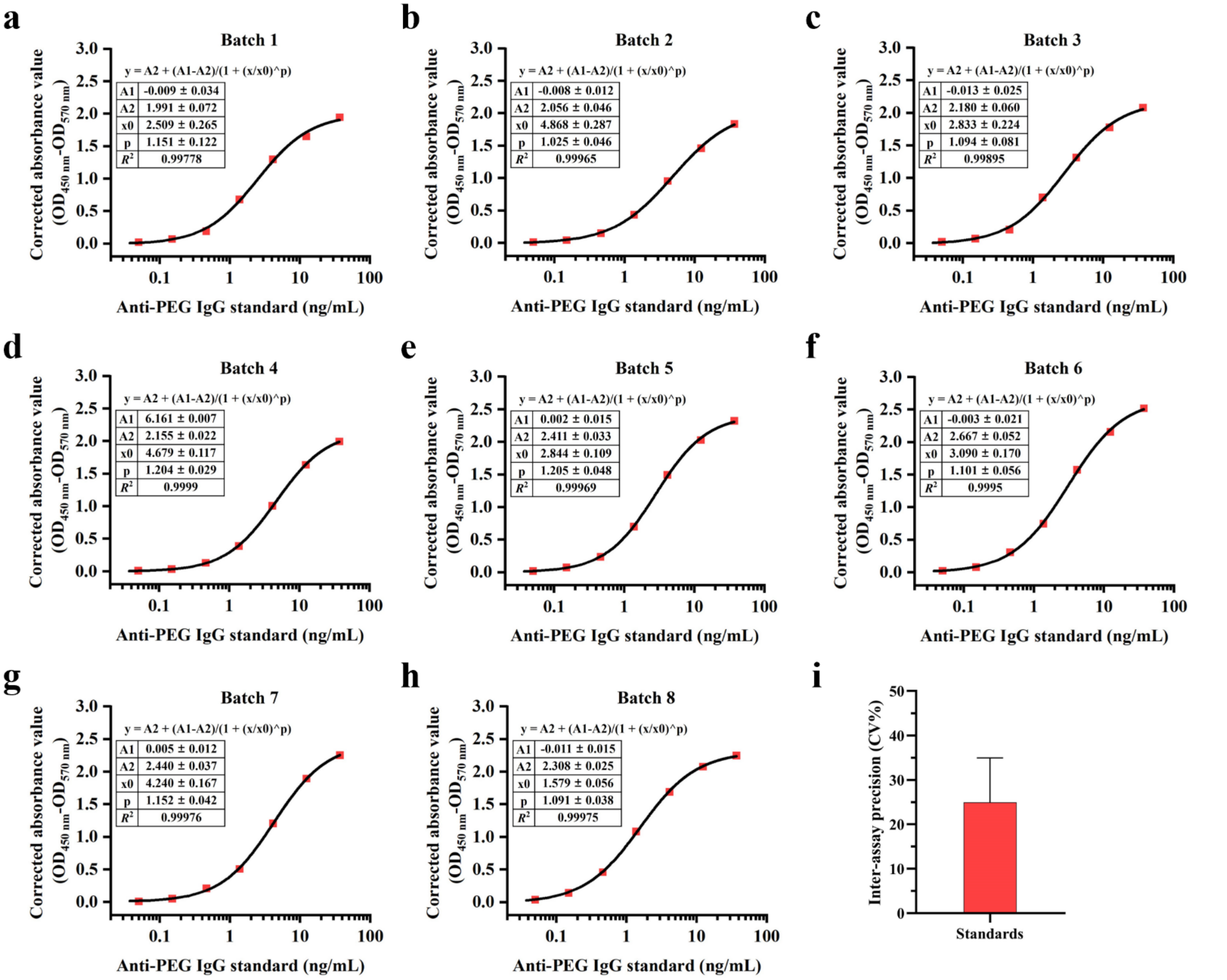


**Supplementary Fig. 6. Standard curves of ELISA for detecting anti-PEG IgG in rat serum samples (a-h) and inter-assay precision (CV%) of anti-PEG IgG standards (i).** Standard curves were constructed by plotting the average absorbance values ($OD_{450\ nm}$-$OD_{570\ nm}$) and corresponding antibody concentrations with Four Parameter Logistic (4PL) curve fit using Origin 2021 software. Serial dilutions of anti-PEG IgG standards (0.05, 0.15, 0.46, 1.37, 4.12, 12.35, 37.04 ng/mL) were included in each batch of ELISA for total eight independent batches. Inter-assay precision was determined by calculating the Coefficient of Variation (CV% = (Standard deviation/Mean) ×100%) for anti-PEG IgG standards among all eight batches of ELISA, which was 24.896 ± 10.071% as indicated in subfigure **i** (see **Methods** for acceptance criteria). In addition to the anti-PEG IgG standards run for each batch, 88 different rat serum samples were respectively tested in batch 1-3, 5 and 7, and 54 different rat serum samples were respectively tested in batch 4. In batch 6 and 8, 71 different rat serum samples were respectively tested. Data in **i** were presented as “mean ± standard deviation” (n = 7).

**Supplementary Table 3. Linear mixed model analysis of change in the anti-PEG IgM level after injection of PEGylated LNP over time across groups**

| Variable | Anti-PEG IgM | | | | | |
|---|---|---|---|---|---|---|
| | β (95% CI) | *P* | β (95% CI) | *P* | β (95% CI) | *P* |
| **Group** | | | | | | |
| Control | 0 (ref.) | - | - | - | - | - |
| L-LNP | 0.2337 (-0.0351, 0.5025) | 0.0915 | 0 (ref.) | - | - | - |
| M-LNP | 0.6198 (0.3509, 0.8886) | <0.0001 | 0.3861 (0.1618, 0.6103) | 0.0011 | 0 (ref.) | - |
| H-LNP | 0.4103 (0.1415, 0.6792) | 0.0035 | 0.1767 (-0.0476, 0.4009) | 0.1257 | -0.2094 (-0.4336, 0.0148) | 0.0701 |
| **Time** | 0.0140 (0.0032, 0.0249) | 0.0116 | - | - | - | - |
| **Time$^2$** | -0.0008 (-0.0009, -0.0006) | <0.0001 | - | - | - | - |
| **Group*Time** | | | | | | |
| Control*Time | 0 (ref.) | - | - | - | - | - |
| L-LNP*Time | 0.0034 (-0.0036, 0.0105) | 0.3408 | 0 (ref.) | - | - | - |
| M-LNP*Time | 0.0238 (0.0167, 0.0308) | <0.0001 | 0.0203 (0.0145, 0.0262) | <0.0001 | 0 (ref.) | - |
| H-LNP*Time | 0.0458 (0.0387, 0.0528) | <0.0001 | 0.0424 (0.0365, 0.0482) | <0.0001 | 0.0220 (0.0161, 0.0279) | <0.0001 |
| **Injection** | | | | | | |
| First | 0 (ref.) | - | - | - | - | - |
| Second | 0.9166 (0.7852, 1.0479) | <0.0001 | - | - | - | - |

Models considered variables including group, time, time$^2$, number of injections, and interaction term of group and time as fixed effect and subject as random effect. β for group represents mean differences in antibody levels between groups at all time points. β for time and time$^2$ represents rate of change in antibody levels over time for the four groups at all time points. β for injection represents mean difference in antibody levels for the four groups at all time points between the first injection and second injection. β for group*time represents mean differences in the rate of change of antibody levels over time between groups. ref: reference.

**Supplementary Table 4. Linear mixed model analysis of change in the anti-PEG IgG level after injection of PEGylated LNP over time across groups**

| Variable | Anti-PEG IgG | | | | | |
|---|---|---|---|---|---|---|
| | β (95% CI) | *P* | β (95% CI) | *P* | β (95% CI) | *P* |
| **Group** | | | | | | |
| Control | 0 (ref.) | - | - | - | - | - |
| L-LNP | -0.0950 (-0.2748, 0.0848) | 0.3033 | 0 (ref.) | - | - | - |
| M-LNP | 0.0871 (-0.0927, 0.2669) | 0.3449 | 0.1821 (0.0321, 0.332) | 0.0195 | 0 (ref.) | - |
| H-LNP | 0.1230 (-0.0568, 0.3028) | 0.1835 | 0.2179 (0.068, 0.3679) | 0.0054 | 0.0359 (-0.1141, 0.1858) | 0.6404 |
| **Time** | -0.0092 (-0.0159, -0.0024) | 0.0077 | - | - | - | - |
| **Time$^2$** | -0.0001 (-0.0002, -0.00002) | 0.0197 | - | - | - | - |
| **Group*Time** | | | | | | |
| Control*Time | 0 (ref.) | - | - | - | - | - |
| L-LNP*Time | 0.0011 (-0.0033, 0.0054) | 0.6339 | 0 (ref.) | - | - | - |
| M-LNP*Time | 0.0149 (0.0105, 0.0193) | < 0.0001 | 0.0138 (0.0102, 0.0175) | < 0.0001 | 0 (ref.) | - |
| H-LNP*Time | 0.0244 (0.0200, 0.0288) | < 0.0001 | 0.0233 (0.0197, 0.027) | < 0.0001 | 0.0095 (0.0059, 0.0131) | < 0.0001 |
| **Injection** | | | | | | |
| First | 0 (ref.) | - | - | - | - | - |
| Second | 0.6549 (0.5734, 0.7364) | < 0.0001 | - | - | - | - |

Models considered variables including group, time, time$^2$, number of injections, and interaction term of group and time as fixed effect and subject as random effect. β for group represents mean differences in antibody levels between groups at all time points. β for time and time$^2$ represents rate of change in antibody levels over time for the four groups at all time points. β for injection represents mean difference in antibody levels for the four groups at all time points between the first injection and second injection. β for group*time represents mean differences in the rate of change of antibody levels over time between groups. ref: reference.

**Supplementary Table 5. Linear mixed model analysis of change in the difference of anti-PEG IgM level (▲Anti-PEG IgM ($Log_{10}$ CONC)) between two injections of PEGylated LNP over time across groups**

| **Variable** | **▲Anti-PEG IgM ($Log_{10}$ CONC)** | | | | | |
|---|---|---|---|---|---|---|
| | **β (95% CI)** | ***P*** | **β (95% CI)** | ***P*** | **β (95% CI)** | ***P*** |
| **Group** | | | | | | |
| Control | 0 (ref.) | - | - | - | - | - |
| L-LNP | 0.2281 (-0.0915, 0.5477) | 0.1653 | 0 (ref.) | - | - | - |
| M-LNP | 1.1623 (0.8427, 1.4819) | < 0.0001 | 0.9343 (0.6677, 1.2008) | < 0.0001 | 0 (ref.) | - |
| H-LNP | 1.6775 (1.3579, 1.9971) | < 0.0001 | 1.4494 (1.1828, 1.716) | < 0.0001 | 0.5152 (0.2486, 0.7817) | 0.0003 |
| **Time** | 0.0725 (0.0441, 0.1009) | < 0.0001 | - | - | - | - |
| **Time$^2$** | -0.0026 (-0.0037, -0.0015) | < 0.0001 | - | - | - | - |
| **Group*Time** | | | | | | |
| Control*Time | 0 (ref.) | - | - | - | - | - |
| L-LNP*Time | 0.0045 (-0.0156, 0.0247) | 0.6596 | 0 (ref.) | - | - | - |
| M-LNP*Time | -0.0167 (-0.0369, 0.0034) | 0.1048 | -0.0213 (-0.0381, -0.0045) | 0.0138 | 0 (ref.) | - |
| H-LNP*Time | -0.0165 (-0.0367, 0.0037) | 0.1099 | -0.021 (-0.0379, -0.0042) | 0.0149 | 0.0002 (-0.0166, 0.0171) | 0.9775 |

Models considered variables including group, time, time$^2$, and interaction term of group and time as fixed effect and subject as random effect. β for group represents mean differences in the average levels of ▲Anti-PEG IgM ($Log_{10}$ CONC) among groups at all time points. β for time and time$^2$ represents change rate in ▲Anti-PEG IgM ($Log_{10}$ CONC) over time for the four groups at all time points. β for group*time represents mean differences in the change rates of ▲Anti-PEG IgM ($Log_{10}$ CONC) over time between groups. ▲Anti-PEG IgM ($Log_{10}$ CONC) was defined as Anti-PEG IgM ($Log_{10}$ $CONC_{2nd\ injection}$) ($log_{10}$-transformed concentration of anti-PEG IgM induced during the second injection cycle) subtracting corresponding Anti-PEG IgM ($Log_{10}$ $CONC_{1st\ injection}$) ($log_{10}$-transformed concentrations of anti-PEG IgM induced during the first injection cycle). ref: reference.

**Supplementary Table 6. Linear mixed model analysis of change in the difference of anti-PEG IgG level (▲Anti-PEG IgG ($Log_{10}$ CONC)) between two injections of PEGylated LNP over time across groups**

| Variable | ▲Anti-PEG IgG ($Log_{10}$ CONC) | | | | | |
|---|---|---|---|---|---|---|
| | β (95% CI) | *P* | β (95% CI) | *P* | β (95% CI) | *P* |
| **Group** | | | | | | |
| Control | 0 (ref.) | - | - | - | - | - |
| L-LNP | 0.0149 (-0.1866, 0.2164) | 0.8847 | 0 (ref.) | - | - | - |
| M-LNP | 0.5180 (0.3165, 0.7195) | < 0.0001 | 0.503 (0.335, 0.6711) | < 0.0001 | 0 (ref.) | - |
| H-LNP | 0.8861 (0.6846, 1.0876) | < 0.0001 | 0.8711 (0.7031, 1.0392) | < 0.0001 | 0.3681 (0.2, 0.5362) | < 0.0001 |
| **Time** | 0.1232 (0.1022, 0.1442) | < 0.0001 | - | - | - | - |
| **Time$^2$** | -0.0050 (-0.0058, -0.0042) | < 0.0001 | - | - | - | - |
| **Group*Time** | | | | | | |
| Control*Time | 0 (ref.) | - | - | - | - | - |
| L-LNP*Time | 0.0031 (-0.0118, 0.0180) | 0.6848 | 0 (ref.) | - | - | - |
| M-LNP*Time | 0.0030 (-0.0119, 0.0179) | 0.6899 | -0.0001 (-0.0125, 0.0124) | 0.9933 | 0 (ref.) | - |
| H-LNP*Time | 0.0002 (-0.0147, 0.0151) | 0.9832 | -0.0029 (-0.0154, 0.0095) | 0.6445 | -0.0029 (-0.0153, 0.0096) | 0.6505 |

Models considered variables including group, time, time$^2$, and interaction term of group and time as fixed effect and subject as random effect. β for group represents mean differences in the average levels of ▲Anti-PEG IgG ($Log_{10}$ CONC) among groups at all time points. β for time and time$^2$ represents change rate in ▲Anti-PEG IgG ($Log_{10}$ CONC) over time for the four groups at all time points. β for group*time represents mean differences in the change rates of ▲Anti-PEG IgG ($Log_{10}$ CONC) over time between groups. ▲Anti-PEG IgG ($Log_{10}$ CONC) was defined as Anti-PEG IgG ($Log_{10}$ $CONC_{2nd\ injection}$) ($log_{10}$-transformed concentration of anti-PEG IgG induced during the second injection cycle) subtracting corresponding Anti-PEG IgG ($Log_{10}$ $CONC_{1st\ injection}$) ($log_{10}$-transformed concentrations of anti-PEG IgG induced during the first injection cycle). ref: reference.

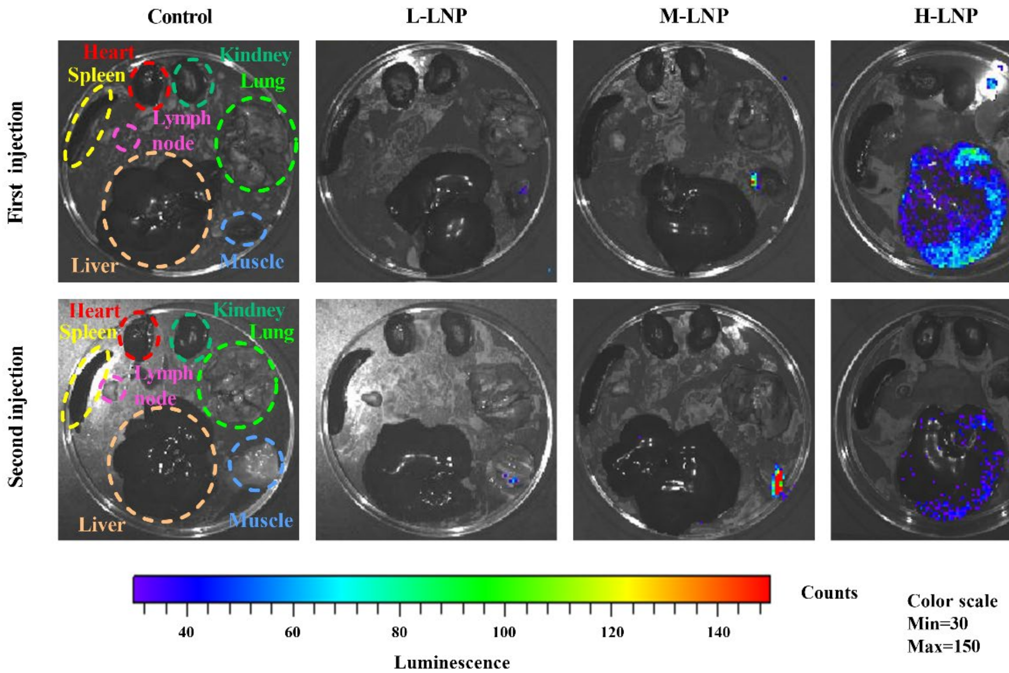


**Supplementary Fig. 7. Representative luminescence images of major organs and muscle tissues isolated from rats 6 hours after the first and second injections of DiR-LU@LNP.** Wistar rats were injected intramuscularly with 0.009 (L-LNP group), 0.342 (M-LNP group) and 2.358 (H-LNP group) mg phospholipids/kg DiR-LU@LNP on Day 0 and Day 21, respectively. Rats in the Control group were injected with PBS. Six hours after each injection, three rats from each experimental group were administered with D-luciferin at a dose of 150 mg/kg intraperitoneally. Fifteen minutes after administration of D-luciferin, rats were sacrificed and immediately dissected. Major organs including heart, liver, spleen, lung, kidneys and draining lymph node, and muscle at the injection site were collected for bioluminescence imaging with IVIS Spectrum imaging system.

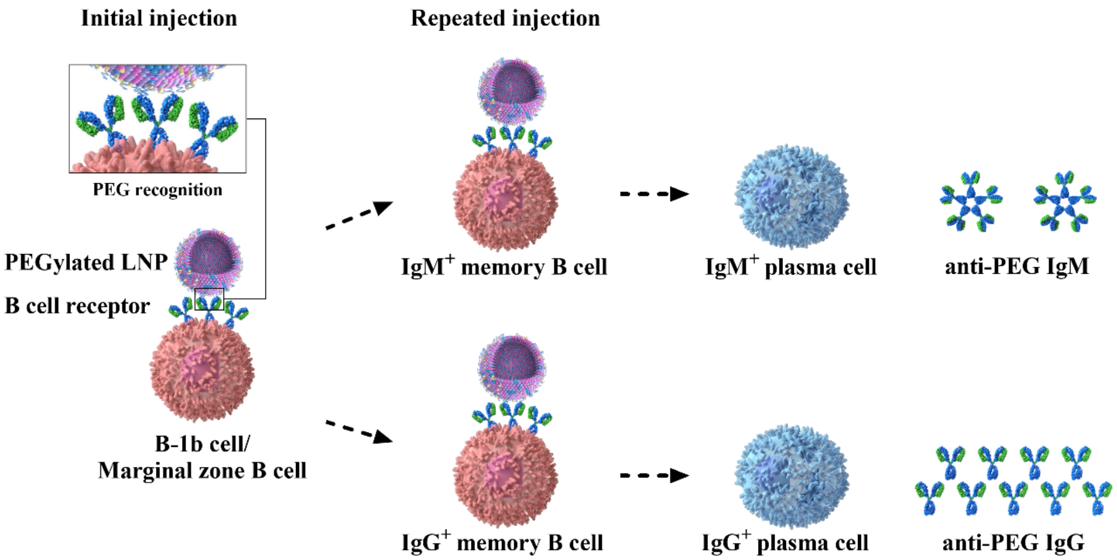


**Supplementary Fig. 8. Hypothetical mechanism for B cell memory induced by PEGylated LNP.** After initial injection of PEGylated LNP, PEG on the surface of LNP extensively cross-links B cell receptors (BCRs), and thereby activate B-1b cells and marginal zone B cells. Following activation, these cells can differentiate into $IgM^+$ memory B cells and $IgG^+$ memory B cells. After repeated injection of PEGylated LNP, pre-existing $IgM^+$ memory B cells and $IgG^+$ memory B cells immediately recognize PEG on the surface of newly injected LNP though BCRs and differentiate into $IgM^+$ plasma cells and $IgG^+$ plasma cells, leading to rapid and intense secretion of anti-PEG IgM and anti-PEG IgG.
